# Estrogen receptor-positive ILC cell line xenografts recapitulate metastatic dissemination and endocrine response of invasive lobular carcinoma

**DOI:** 10.64898/2026.03.17.712396

**Authors:** Nilgun Tasdemir, Laura Savariau, Julie Scott, Joseph D Latoche, Kyle Biery, Zheqi Li, Emily A Bossart, Sreeja Sreekumar, Daniel D Brown, Sarah Wang, Rebecca J Watters, Azadeh Nasrazadani, Ye Qin, Ye Cao, Fangyuan Chen, Matthew J. Sikora, George Tseng, Carlos Castro, Carolyn J Anderson, Jennifer Atkinson, Jagmohan Hooda, Peter C Lucas, Nancy E Davidson, Adrian V Lee, Steffi Oesterreich

## Abstract

**Highlights:**

- ER+, *CDH1*-deficient ILC xenografts recapitulate single-file histology in vivo
- Spontaneous metastases faithfully mirror unique ILC clinical dissemination patterns
- ER+ leptomeningeal metastases confirmed histologically in ILC xenograft models
- Fulvestrant inhibits ILC tumor growth and metastatic burden, improving survival
- ER signaling is enriched in ILC brain metastases, with upregulation of *RET*

Invasive lobular breast carcinoma (ILC), the most common special histological subtype of breast cancer, is characterized by nearly universal expression of estrogen receptor alpha (ER) and unique sites of metastases, neither of which is fully recapitulated by genetically engineered mouse models. Using reporter-labeled ILC mouse xenografts, herein we used mammary fat pad, tail vein and intracardiac orthotopic growth to analyze spontaneous and experimental metastasis and gene expression. We observed ER-positive primary tumors with single-file histology and collagen deposition, and spontaneous metastasis from the mammary fat pad to bones, ovaries, and brain including the leptomeninges, thereby closely mirroring the growth and metastatic spread of human ILC. Brain metastases showed strong ER staining, confirmed by sequencing analyses which identified estrogen signaling as top activated pathway, and the lesions exhibited robust response to endocrine therapy. In summary, we report endocrine responsive mammary fat pad, tail vein and intracardiac xenografts that faithfully demonstrate unique ILC features and can serve as invaluable pre-clinical translational platforms for validating candidate ILC genetic drivers and testing novel therapeutics.

## Introduction

Invasive lobular carcinoma (ILC) is the most common special histological subtype of breast cancer accounting for 10-15% of all breast cancer cases ^1^ and the 6^th^ most common cancer type affecting women in the US ^2^. ILC tumors are characterized by genetic loss of (*CDH1*) E-cadherin ^3^, which disrupts adherens junctions and causes relocalization of p120-catenin from the cell membrane to the cytoplasm ^4,5^, leading to its unique diffuse single-file histology ^6,7^. ILC tumors largely display high expression of estrogen receptor alpha (ER) and progesterone receptor (PR) and a low proliferative index and are treated with endocrine therapy as standard of care ^8,9^. Despite these favorable clinical characteristics, patients with ILC often present with endocrine resistance and late recurrences ^10–12^. In addition to shared sites of metastases with IDC patients such as bone, brain and lung, patients with ILC also exhibit dissemination to unusual anatomical sites including ovary, eye, peritoneum and gastrointestinal tract ^1,13–15^.

To better understand the unique histological and clinical features of ILC, several mouse models have been developed attempting to recapitulate it in the laboratory. The first ILC mouse model was based on combined genetic inactivation of *Cdh1* and *Trp53*, and it captured ILC’s unique histology and dissemination ^16^. Multiomics profiling of ILC ^3,17,18^ has identified activation of the PI3K pathway as a major driver of ILC, and mouse models have been developed modeling this pathway by combining loss of *Cdh1* with i) loss of *Pten* ^19^, and ii) with activating *Pik3ca* mutations ^20^. While these genetically engineered mouse models have increased our understanding of ILC biology, they have not fully recapitulated all unique clinical features such as ovarian metastases and robust estrogen response. For a comprehensive review of available ILC experimental models including cell lines, organoids, genetically engineered mouse models, and patient-derived xenografts, see Sflomos et al. ^21^ More recently, ILC cell line xenograft models have been developed via intraductal injections into immunocompromised mice, with reports of metastases to sites such as ovary ^22,23^. Here, we set out to complement these approaches through systematic development and characterization of comprehensive xenograft models using a larger set of human ILC cell lines injected via mammary fat pads and assessment of experimental metastasis capacity.

We previously reported an extensive *in vitro* characterization of ER-positive human ILC cell lines, which exhibited differential hormone responses ^24,25^, as well as unique growth and migration characteristics and gene expression patterns as compared to human IDC cell lines ^26,27^. Here, we extend those findings to *in vivo* growth and dissemination features, endocrine response and transcriptional profiling. We find that mammary fat injections of human ILC cell lines give rise to tumors closely recapitulating the slow growth, hormone receptor expression and discohesive growth of human ILC tumors within dense stroma. Both spontaneous metastases arising from orthotopic injections and experimental metastases reflect clinical patterns of ILC dissemination to bone, ovary, and brain, including leptomeningeal metastases and other sites. The metastatic lesions responded to endocrine therapy, in line with results from transcriptional profiling of primary tumors and spontaneous metastases to the brain that revealed an enrichment for ER signaling in brain metastases. Collectively, our comprehensive study augments our current understanding of unique ILC biology and existing *in vivo* models available to the global research community, which should better enable efforts to improve the clinical management and outcome of patients with ILC.

## Materials and Methods

### Cell culture

MDA-MB-134-VI (MDA-MB-134) and MDA-MB-330 were obtained from the American Type Culture Collection (ATCC). SUM44PE (also abbreviated as SUM44) was purchased from Asterand. BCK4 cells were kindly provided by Britta Jacobsen, University of Colorado Anschutz, CO, under a Material Transfer Agreement that precludes the authors from sharing the cell line. Any requests for the BCK4 cells should be directed to the University of Colorado Anschutz. Cell lines were maintained as previously described ^26^ in the following media (Life Technologies) with 10% FBS: MDA-MB-134 and MDA-MB-330 in 1:1 DMEM:L-15, BCK4 in MEM with non-essential aminoacids (Life Technologies) and insulin (Sigma-Aldrich). SUM44 was maintained in DMEM-F12 with 2% charcoal stripped serum and supplements. Cell lines were routinely tested to be mycoplasma free, authenticated by the University of Arizona Genetics Core by Short Tandem Repeat DNA profiling and kept in continuous culture for <6 months. Cells were infected with lentivirus encoding Luciferase and tRFP (pLEX-TRC210/L2N-TurboRFP-c ^28^) as previously described ^29^, selected with neomycin and FACS-sorted for the top 5% brightest tRFP expression.

### Animal experiments

Animal experiments were performed according to the University of Pittsburgh Institutional Animal Care and Use Committee (IACUC) guidelines. 5–6-week-old female NOD-scid IL2Rgamma null (NSG) mice (JAX stock #: 005557) and 3–4-week-old athymic nude (Foxn1^nu^) mice were obtained from The Jackson Laboratory (JAX stock #: 002019). Mice were allowed to acclimate for two weeks before use for experiments. For mammary fat pad injections, 5,000,000 cells were injected into the 4^th^ inguinal glands of 7–8-week-old NSG mice in 1:1 serum free media and phenol-red free Matrigel (Corning, #356237) using a 26G syringe in 50 μl volume total. For tail vein injections, 5,000,000 MDA-MB-134/SUM44 and 2,500,000 BCK4/MDA-MB-330 cells were injected into the lateral tail vein of 7–8-week-old nude or NSG mice in 100ul of PBS under a heat lamp. For intracardiac injections, 150,000-500,000 cells were injected into the left ventricle of 5–6-week-old nude mice in 100 μl of PBS under isoflurane anesthesia (Patterson Veterinary, #78938441). Mice were implanted with 0.36mg 90-day release estradiol pellets (Innovative Research of America) or with ∼0.4mg homemade estradiol pellets with beeswax. For primary tumor removal experiments, tumors of 300-800mm^3^ volume were excised by performing survival surgery on mice under sterile conditions and anesthesia. To combat estradiol supplementation-related infections, mice were prophylactically treated with 0.1mg/ml Baytril (Valley Vet Supply; # 328RX).

Tumor volumes were monitored weekly by caliper measurements. Tumor volumes were calculated using the formula V=LxW^2^/2. *In vivo* and *ex vivo* bioluminescent imaging was performed using IVIS200 and IVIS LuminaXR imaging systems (PerkinElmer) following intraperitoneal injection of 150 mg/kg D-luciferin (Gold Bio, #LUCK-100). For *in vivo* bioluminescent imaging, both dorsal and ventral images were captured. For orthotopic injections, primary tumors were covered with a piece of black paper to block the saturating signal to visualize the less intense metastatic signals. For *ex vivo* bioluminescent imaging, mice were sacrificed 5 minutes after luciferin injection and organs were quickly harvested and imaged in 6-well plates in PBS. For visualizing tRFP expression *ex vivo,* images were captured under a fluorescent stereomicroscope (Olympus SZX16) immediately following harvesting of mice and dissection of organs. For the fulvestrant experiments, when the mammary fat pad tumors reached 100-200mm^3^ or intracardiac metastatic signal total flux reached 0.5-1E^8^, mice were randomized to sub-cutaneous treatment in the nape of the neck with vehicle (peanut oil) or 5mg fulvestrant (Selleck Chemicals, #S1191) bi-weekly.

To further study leptomeningeal metastases formation, 5×10^6^ SUM44PE and MDA-MB-134 cells were orthotopically injected into the mammary fat pad of 10 NSG mice for each cell line. Primary tumors were surgically excised when they reached a volume of 200 mm³. Post-surgery, tumor progression was monitored weekly using *in vivo* IVIS imaging. For each imaging session, 100 ul of luciferase substrate was injected intraperitoneally, and imaging was performed 5 minutes later. To enhance visualization of brain metastasis, the body of each mouse was shielded to minimize luminescence from other tissues.

The first detectable brain metastasis signal on IVIS scans was designated as “baseline” for subsequent staging. Given the variability in tumor growth and metastasis among individual mice, brain metastasis progression was assessed based on luminescent signal intensity. One mouse in the MDA-MB-134 group and four in the SUM44PE group died before exhibiting brain metastasis. For the remaining mice, approximately half of the mice in each group exhibited strong brain signals, while the other half showed weaker luminescence signals. These differences were used to define “early” and “late” stages of brain metastasis., this divergence in signal intensity occurred at ∼140 days post tumor cell injection for the SUM44PE group and ∼120 days post tumor cell injection for the MDA-MB-134 group. At these respective time points, all mice within each group were euthanized for tissue collection and analysis. Brain tissues were collected and examined for RFP signals under a dissection microscope. The tissue samples were then fixed in formalin and embedded in paraffin (FFPE), followed by immunohistochemical staining with human cytokeratin-19 (Fisher Scientific #MS198P1) to evaluate leptomeningeal metastasis.

### Tissue processing and staining

Tissues were fixed overnight in 10% formalin solution (Sigma-Aldrich #HT501128) and then switched to 70% ethanol. Bone samples were decalcified for 7-10 days using Osteosoft (Millipore Sigma, #101728), with buffer changes occurring every 2 days. Fixed/decalcified tissues were embedded into paraffin and sectioned onto slides at 4 microns. ERα, PR, CDH1 (Ecad) and p120 staining was performed on a Ventana automated immunostainer. For PDGFRβ and Ly6G staining, the slides were baked for at least 1 hour at 65°C, deparaffinized and rehydrated by immersing in xylene and a series of decreasing alcohol concentrations. Antigen retrieval was performed in Tris EDTA (pH9) for PDGFRβ and citrate buffer (pH6) for Ly6G in a pressure cooker for 20 minutes. The slides were treated with hydrogen peroxidase (Abcam #ab64218) for 5 min to block endogenous peroxidase activity. The slides were rinsed with PBS-T (0.5% Tween20) and incubated for 1 hour with 3% Bovine Serum Albumin blocking buffer (Sigma-Aldrich #A9647) in PBS-T. Slides were incubated overnight at 4°C with primary antibody diluted in blocking buffer. Primary antibodies were as follows: ERα (clone SP1 from Roche, #05278414001), CDH1 (Clone 36 from Roche, #05905290001), p120 (clone 98/pp120, BD Biosciences, BD610134), PR (clone 1E2, Roche, 05278392001), PDGFR (Cell Signaling Technologies, CST #3169, 1:100), and Ly6G (BD, #551459, 1:100). The next day, slides were rinsed 3 times in PBS-T before a 45min incubation with appropriate secondary antibodies (Mouse DAKO – K4001, Rabbit DAKO – K4011). The slides were rinsed with PBS-T followed by staining with DAB (Agilent K3468). The slides were counterstained in hematoxylin before rinsing in water and dehydration by immersing in a series of increasing alcohol concentrations and xylene. Trichrome staining was performed using the Trichrome Stain (Masson) kit (Sigma Aldrich, #HT15-1KT) according to the manufacturer protocols. Before imaging, the slides were mounted with permount (Fisher Scientific #SP15100).

### RNA sequencing

To enrich for disseminated tumor cells at these secondary sites, the metastatic lesions were macro-dissected *ex vivo* by visualizing the tRFP expression under a fluorescent stereomicroscope (Olympus SZX16). Tissues were stored in RNAlater at −80^0^C. RNA extraction was performed as previously described ^30^. Small pieces of tissue were cut out and weighed to not use more than 30mg and overload the columns. They were put in white-capped FACS tubes in 600 μl Buffer RLT+BME. The hand-held Homogenizer 150 (Fisher Scientific) was used with plastic probes, which were extensively cleaned with 70% EtOH in between samples. Then the samples were run through the QIAShredder for further homogenization. RNA was isolated using the Qiagen RNeasy Mini Kit protocol and on column DNase digestion.

RNA concentrations were quantified by ThermoFisher Qubit (Cat # Q33238) and their size distribution measured by Agilent TapeStation (Cat # G2991AA). Library prep and mRNA sequencing was performed by the Pitt Genomics Core. Briefly, paired-end RNAseq libraries were prepared using KAPA poly A RNA hyperPrep kit (Cat # 8098093702), and sequenced with NovaSeq 6000 for a target of 100 million reads/sample using S1 reagents for 300 cycles, 0.8 billion reads and total output of 250G (Cat # 20028317). FastQ files were checked for quality by MulitQC (1.8) before mapping to the Ensembl GRCh38 reference genome with STAR (2.7.5a). Subsequently, BAM files were filtered with XenofilteR (1.6) to separate the human reads from the cell line xenograft and from the reads from surrounding mouse tissue. Both human and mice reads were recovered. Finally, reads were quantified with featureCounts using reference Homo.sapiens.GRCh38.100 for human reads and Mus.musculus.GRCm39.100. Tximport (1.16.1) was used to generate final matrix counts in transcript per million (TPM) which was subsequently used in R for analysis. Reads were mapped to reference human and mouse transcriptomes and XenoFilter algorithm was applied.

DESeq2 (1.30.0) in R (4.0.0) was used to identify differential gene expression (DGEs). Tidyr (1.1.3) was used to re-arrange the matrix with the cell line and cancerous lesions to first contrast the cell line vs. primary tumor followed by primary tumors vs. brain metastases and primary tumor vs. ovarian metastases. PCA plots were generated with plotPCA function of DESeq2 and visualized with ggplot2 (3.3.5). Volcano plots were generated with EnhancedVolcano (1.6.0) using genes from the DGE analyses. GSEA was performed with ClusterProfiler (3.16.1) and tested for all gene sets. The top 15 most significant pathways identified were graphically represented using ggplot2.

### Circulating free DNA (cfDNA) extraction and digital droplet PCR (ddPCR)

cfDNA extraction and ddPCR were performed as previously described ^31^. In brief, mouse blood samples were collected in EDTA tubes (BD, #367856) and cfDNA was isolated from plasma samples using Qiagen circulating nucleic acid kit (#55114). KRAS and MAP2K4 genes were pre-amplified in cfDNA using primers: KRAS G12 - TAAGCGTCGATGGAGGAGT (F) and GTATCAAAGAATGGTCCTGCACC (R), and MAP2K4_S184 – CCTGTCCCTGTCCATACAATTA (F) and GAACATTGGCAGTATTTATCGAGAC (R) with GoTaq Mastermix system and 40 cycles of amplification. Pre-amplified products were further diluted (10,000x) and subjected for droplet digital PCR detection using QC200X system following manufacturer protocol. Droplets were analyzed using BioRad droplet reader and QuantaSoft software (BioRad, Version 1.7) was used to quantify mutation allele frequency. Specific probes for ddPCR detection were used: MAP2K4 S184 probe (WT) – TCTACCTCGTTTGAT (FAM), and MAP2K4 L184 probe (Mutant) – TCTACCTTGTTTGAT (VIC); KRAS G12 probe (WT) – GGAGCTGGTGGCGTA (FAM), and KRAS R12 probe (Mutant) – GGAGCTCGTGGCGTA (VIC). Cell-free DNA extracted from plasma of two NSG mice and genomic DNA extracted from MDA-MB-134 cells were used as negative and positive controls, respectively.

### Ex vivo micro computed tomography (μCT)

High resolution CT imaging of non-decalcified, fixed bones was performed using a Siemens Inveon Multimodality scanner, and CTs were 80 kV, high magnification, binning =0, Feldkamp reconstruction with downsample = 0. Scans were approximately 13 minutes each. CT scans were visualized as individual slices or using 3D MIP view and Siemens IRW 3D viewer.

### Statistical analysis

Data analysis and plotting was performed using GraphPad Prism. Data summary is presented as mean +/- standard deviation. Statistical tests used for each figure are indicated in the respective figure legends. The survival plots and log-rank Mantel-Cox test were performed to assess significance. We modeled *in vivo* tumor growth using a linear mixed-effects model with log-transformed tumor volume to compare the growth rate between cell lines, the fitted straight of log-transformed tumor volume against day shows approximately linear trends for each cell lines. To identify changepoints in tumor growth, we analyzed each growth curve using the R package “strucchange”, which detects structural breaks in regression models. The detected changepoint marks the transition from a stable tumor size to the onset of accelerated growth. Prior to analysis, samples with essentially flat curves (no or limited growth) were excluded.

## Results

### ILC cell line orthotopic xenografts generate primary tumors recapitulating human disease

To assess the potential of human ILC cell lines to generate orthotopic tumors in mice, we used the ER-positive human ILC cell lines MDA-MB-134, SUM44, MDA-MB-330 and BCK4. All cell lines exhibit a genetic loss of E-cadherin (*CDH1*), which is a hallmark of ILC. The exception is MDA-MB-330 that contains a biallelic inactivating mutation in *CTNNA1* (alpha catenin)^7,26,32^., an alternative mechanism for loss of adherens junctions. These cells were labeled with a lentiviral reporter that co-expresses luciferase and tRFP to allow *in vivo* and *ex vivo* tracking of tumor growth and dissemination (**Figure 1A**). Following flow sorting for highest tRFP expression, they were injected into the mammary fat pads of immunocompromised NOD-scid IL2Rgamma null (NSG) mice ^33^, which were supplemented with exogenous estradiol and monitored over time.

**Figure 1.**
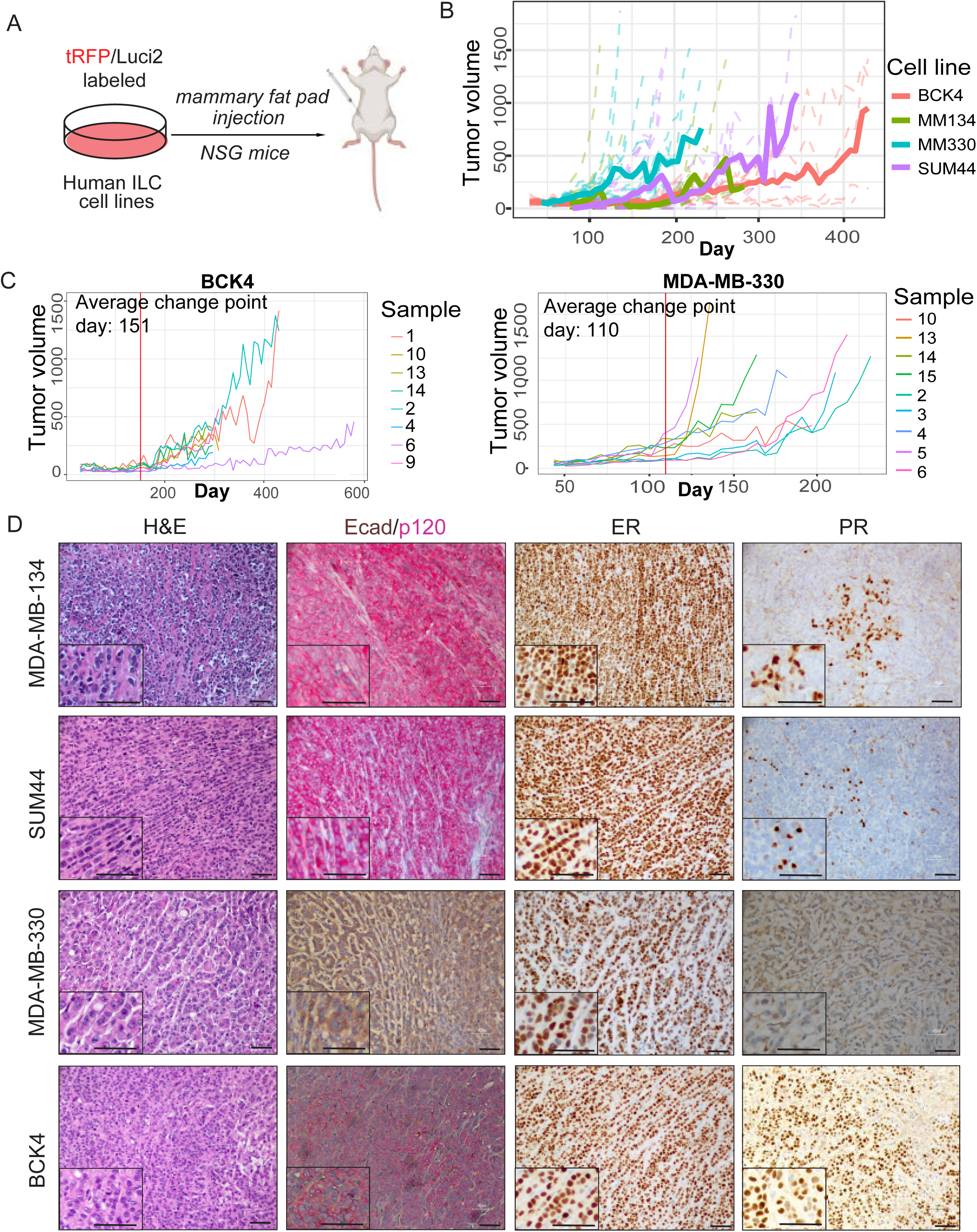
Human ILC cell line orthotopic xenografts generate tumors recapitulating classical ILC histology and ER expression. (A) Schematic of the experiment. (B) Individual tumor volumes from mice with MDA-MB-134 (green), MDA-MB-330 (blue), SUM44PE (purple) and BCK4 (red) orthotopic xenografts. The solid line shows the average for each cell line. N=13 tumors for BCK4 and MDA-MB-134; N=14 tumors for MDA-MB-330; N=15 tumors for SUM44. (C) Quantification of delayed tumor onset via calculation of changepoint in tumor growth curve that detects structural breaks in regression models. Graphical presentation of individual tumor growth curves for BCK4 and MDA-MB-330 with change point indicated in red (days 151 and 110, respectively). (D) H&E, Ecad/p120, ER and PR staining of representative MDA-MB-134, SUM44PE, MDA-MB-330 and BCK4 orthotopic tumors. Insets show a small portion of the images at higher magnification. Scale bar: 40 μm.

Tumor caliper measurements revealed a generally slow *in vivo* growth pattern (**Figure 1B** and **Figure S1A**), consistent with the slow *in vitro* growth rate of these cell lines ^26^ and the slow proliferation rate of human ILC ^3,6^. We modeled the tumor growth using a linear mixed-effects model with log-transformed tumor volume to compare the growth rate between cell lines; the fitted straight of log-transformed tumor volumes against days shows approximately linear trends for each cell line. Results indicated significant differences between most cell lines: MDA-MB-134 grew faster than BCK4 and MDA-MB-330, BCK4 was slower than both MDA-MB-330 and SUM44, while MDA-MB-134 and SUM44 exhibited similar growth rates **(Figure S1B)**. Different growth kinetics was also observed within each cell line, with some tumors initiating early and others displaying a delayed timeline to establish. We did observe consistent delay of tumor growth in BCK4 and SUM44 cells and hence set out to identify changepoints in tumor growth which mark the transition from a stable tumor size to the onset of accelerated growth. Growth curves were analyzed using the R package “strucchange”, which detects structural breaks in regression models. Changepoints for BCK4 and MDA-MB-330 were at day 151 and 110, respectively **(Figure 1C and S1C)**, indicating delayed onset of tumorigenesis in these models.

These reporter-expressing tumors were visualized *in vivo* by bioluminescent imaging (**Figure S1D**) and *ex vivo* by fluorescence stereomicroscopy (**Figure S1E**). Histological characterization by hematoxylin and eosin (H&E) staining revealed a single-file tumor growth pattern for all cell lines (**Figure 1D**), consistent with the classical ILC morphology in human ^6^. Dual immunohistochemistry (IHC) confirmed the genetic/functional loss of E-cadherin and re-localization of p120-catenin from the cell membrane to the cytoplasm in MDA-MB-134, SUM44, and BCK4 cells. Tumors from all cell lines displayed widespread and high levels of ER expression. PR was expressed highest in BCK4, with limited expression in MDA-MB-134 and SUM44 and near absent in MDA-MB-330. These data are consistent with previous reports of *in vitro* ER and PR expression in these cell lines ^26^.

### ILC cell line orthotopic xenografts give rise to spontaneous metastases closely mirroring human disease

In addition to primary tumors, ILC cell line xenografts gave rise to spontaneous metastases from the mammary fat pad over time, as visualized by *in vivo* bioluminescent imaging (**Figure 2A**) and *ex vivo* by fluorescence stereomicroscopy (**Figure 2B**). Time to observation of detectable metastases was 3-4 months in MDA-MB-134, 4-5 months in SUM44, 6-7 months in MDA-MB-330 and over 10 months in BCK4 models. Anatomical sites of metastases included the brain, bones, reproductive tract, and lungs which are common sites in patients with ILC ^1,12^, as well as less common sites such as adrenal glands. These metastatic lesions were confirmed histologically by H&E staining and dual E-cadherin/p120 IHC and appeared largely ER positive (**Figures 2C** and **2D; Figures S2A** and **S2B**), like their respective primary tumors.

**Figure 2.**
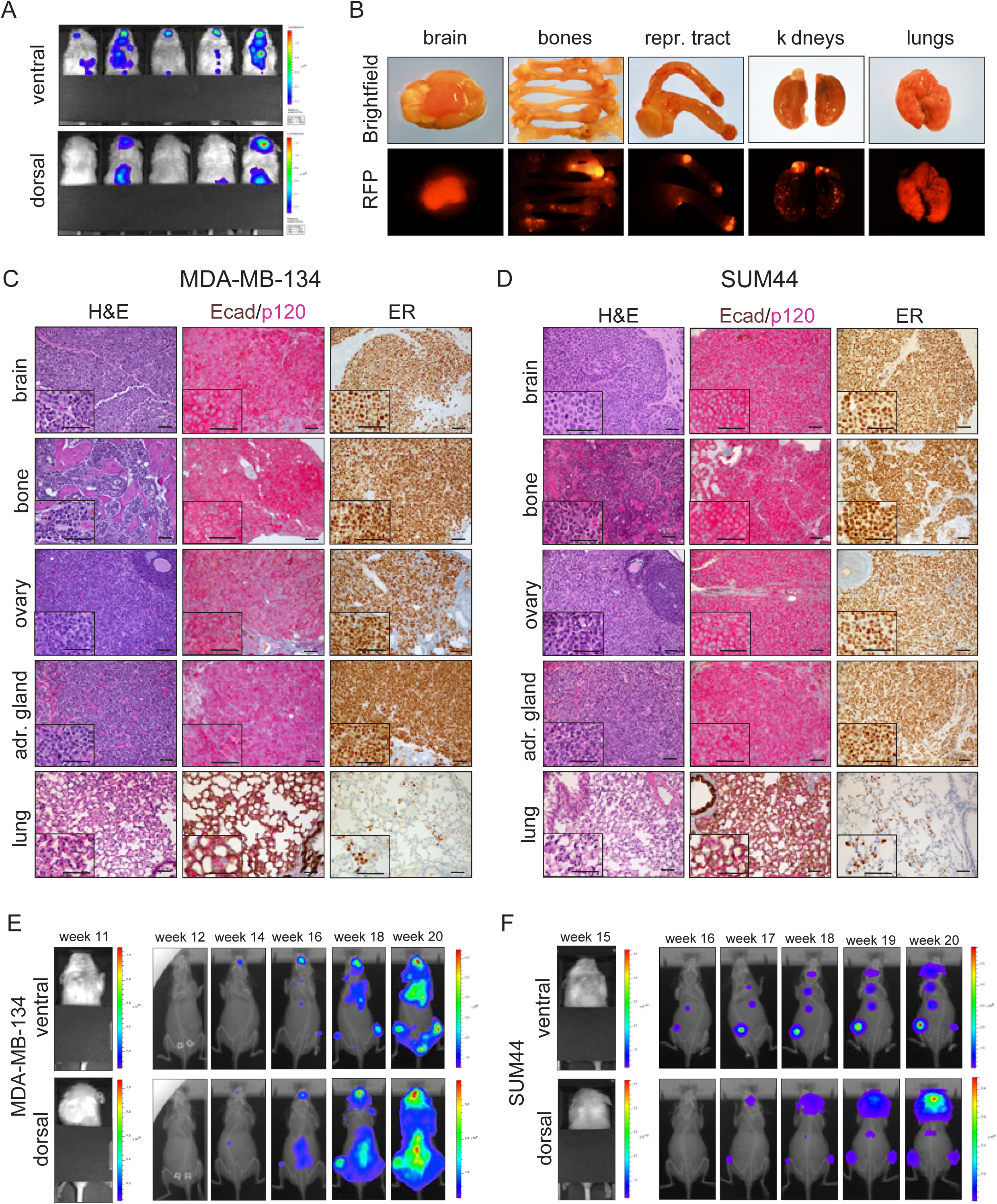
MDA-MB-134 and SUM44PE orthotopic xenografts give rise to spontaneous ER+ metastases to clinically relevant anatomical sites. (A) Ventral (top) and dorsal (bottom) in vivo bioluminescent imaging of mice with MDA-MB-134 orthotopic xenografts showing upper body metastases. The lower bodies have been covered to avoid saturating the camera with the strong signals from the primary tumors – see Figure 1C. (B) Brightfield (top) and RFP (bottom) images of metastases from MDA-MB-134 orthotopic xenografts to the indicated organs. (C-D) H&E, Ecad/p120 and ER staining of representative metastases from MDA-MB-134 (C) and SUM44PE (D) orthotopic xenografts to the indicated organs. Insets show a small portion of the images at higher magnification. Scale bar: 40 μm. (E-F) Ventral (top) and dorsal (bottom) in vivo bioluminescent imaging of mice with MDA-MB-134 (E) and SUM44PE (F) orthotopic xenografts before (left) and for several weeks after (right) removal of primary tumors. In the left panels, the primary tumors have been covered to show the absence of detectable metastases at the time of tumor removal.

While the metastatic lesions to most sites were generally focal, dissemination to the lungs appeared widely scattered (**Figures 2C** and **2D; Figures S2A** and **S2B**). ER-positive reproductive tract metastases were localized to both ovaries (**Figures 2B-2D**) and the uterine horn (**Figures 2B** and **S3A**). ER-positive osseous metastases were seen in long bones (**Figures 2B-2D**) and in the mandible (**Figure S4A and S4B**). High resolution *ex vivo* micro-computed tomography (μCT) revealed signs of bone erosion at the sites of metastases but not in unaffected bones (**Figures S4C-S4F**), consistent with observations of lytic osseous metastases in the clinic ^34,35^.

To determine if the primary tumors needed to be continually present for metastatic dissemination, MDA-MB-134 and SUM44 orthotopic xenografts were established and monitored by bioluminescent imaging. Survival surgery was performed by removal of primary tumors prior to the appearance of detectable metastases, which was at 11 weeks for MDA-MB-134 (**Figure 2E**) and 15 weeks for SUM44 (**Figure 2F**). Over the next few weeks, small focal lesions appeared around the previously characterized anatomical sites such as bones, adrenal gland and brain and continued to grow and increase in signal. These data show that the continuous presence of the primary tumor is not required to observe metastasis and suggest a model of early dissemination (likely with early micro-metastases that are below the limit of detection).

We next focused on the observed central nervous system metastases. Histological examination revealed that metastatic tumor cells were frequently located adjacent to blood vessels within the leptomeningeal lining (**Figure 3A**), a pattern consistent with leptomeningeal metastases (LM) seen in patients with ILC ^36,37^. We confirmed loss of E-cadherin and expression of ER in the LM (**Figure S3B**). To further evaluate the ability of the ILC cell lines to induce LM, we expanded the original study and injected MDA-MB-134 and SUM44PE cells into the mammary fat pads and surgically resected the primary tumors when they reached 200 mm³ (**Figure 3B**). Brain metastases were detected in 9/10 MDA-MB-134-injected mice, and in 5/10 SUM44-injected mice (five died before observation). In the MDA-MB-134 group, mice with early-stage brain metastases (N=4) were euthanized ∼5 weeks post-baseline, while mice with late-stage brain metastases (N=5) were euthanized ∼7 weeks post-baseline, with baseline defined as timepoint of first IVIS signal in brain. In the SUM44 group, mice classified having early-stage (N=3) and late-stage (N=2) brain metastases were euthanized at ∼3 weeks and ∼5 weeks post-baseline, respectively.

**Figure 3.**
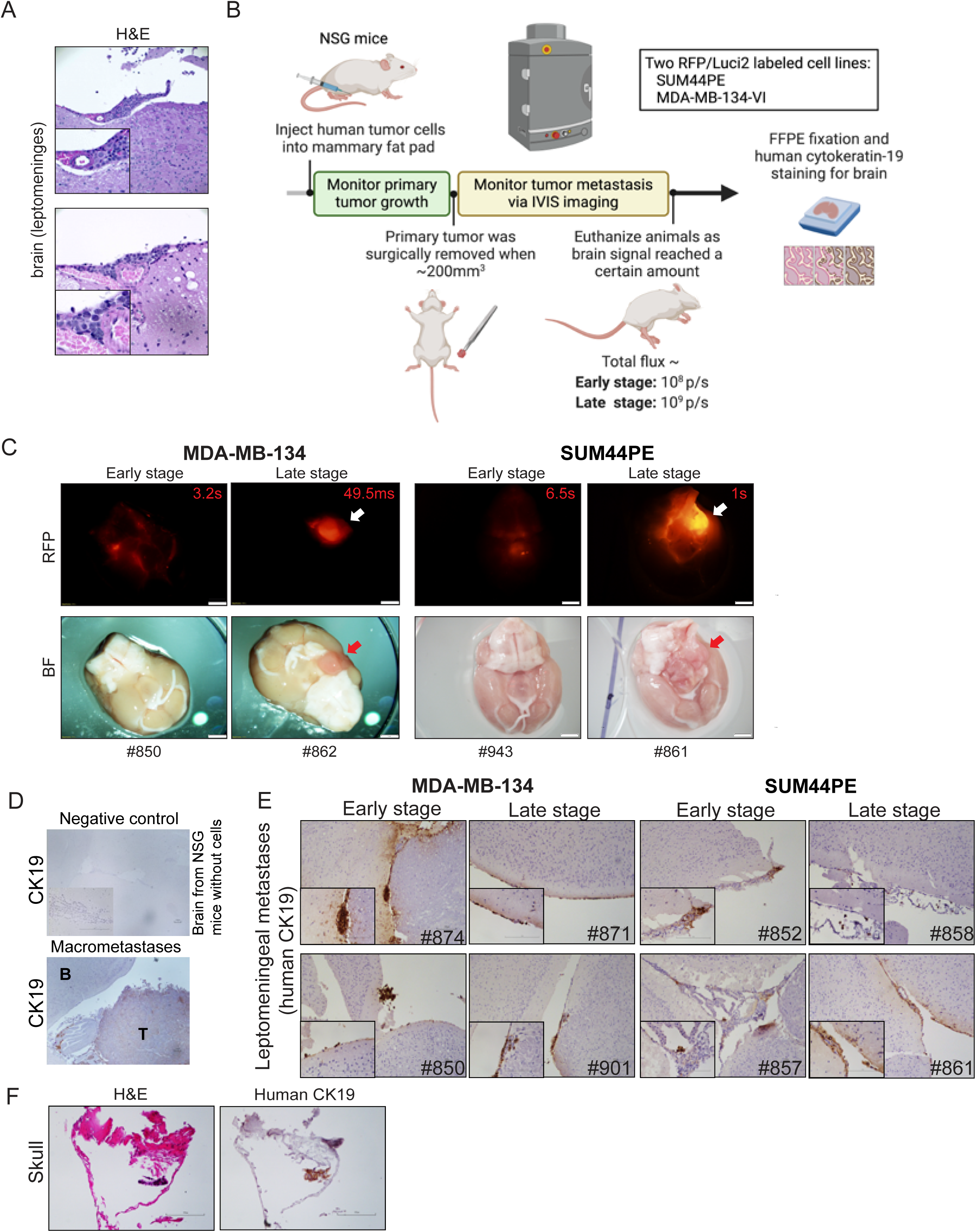
SUM44PE and MDA-MB-134 orthotopic xenografts give rise to leptomeningeal metastases in NSG mice. (A) H&E staining of MDA-MB-134 orthotopic xenograft metastases to the leptomeninges. Scale bar: 40 μm. (B) Schematic representation of the experimental workflow detailing the development of leptomeningeal metastasis through orthotopic injection of SUM44PE and MDA-MB-134 cell lines. (C) Brightfield and RFP images display the ventral sides of mouse brains with either early-stage or late-stage brain metastases, as determined by bioluminescence signal intensity. In late-stage cases, macrometastases are visible on the brain. Scale bar represents 100 μm. (D) IHC with anti-human CK19 antibodies shows leptomeningeal metastases in mice with macrometastases, compared to NSG mice controls without cell injections. B indicates brain regions lacking tumor cells; T marks area with tumor cells. (E) IHC with anti-human CK19 antibodies detected leptomeningeal metastases in mice injected with MDA-MB-134 or SUM44 cells at early and late stages. Shown are images from two mice per group. Scale bar: 100 μm. (F) H&E and CK19 IHC reveal skull micrometastases from late-stage MDA-MB-134 orthotopic injection. Scale bar: 40 μm

In mice with early-stage brain metastases, individual tumor cells were scattered within the leptomeningeal regions without significant macrometastases. In contrast, mice with late-stage metastases exhibited macrometastases predominantly localized to the ventral midbrain near the pons, with additional tumor cells scattered in the dorsal leptomeningeal regions (**Figures 3C**). These tumor cells were either floating in the cerebrospinal fluid or attached to the leptomeningeal tissue. Human origin of the metastatic cells was confirmed using antibodies specific for human CK19 (**Figure 3D&E**). Upon necropsy, visible brain metastases primarily located on the ventral brain surface, were observed in 3/5 MDA-MB-134 mice with late-stage disease, but none with early-stage disease.

Similarly, in the SUM44PE cohort, visible brain macrometastases developed in only 1/3 and 1/2 mice with early-stage and late-stage metastases, respectively. Notably, visible metastatic lesions were observed on the internal surface of the skull in MDA-MB-134 mice (**Figure 3F**), which upon histological examination were confirmed to be located within the leptomeningeal structures. Finally, IHC confirmed that the LM were ER-positive (**Figure S3B**). Overall, these results demonstrate that the tRFP/luciferase expressing SUM44PE and MDA-MB-134 ILC cells induce ER+ leptomeningeal metastases in our NSG mouse model.

### ILC cell line xenografts give rise to experimental metastases via several routes

Next, we determined whether ILC cell line xenografts generate experimental metastases when injected into mice via several routes. To assess the influence of mouse genetic background on experimental metastasis, we performed tail vein and intracardiac injections in both athymic nude and NSG mice. Tail vein injection leads to initial trapping of cells in pulmonary capillaries and subsequent escape into circulation, giving rise to metastases primarily to the lungs but also to other sites ^38,39^. We injected the tRFP and luciferase expressing ILC cell lines into the tail vein of NSG and athymic nude (Foxn1nu) mice, which are deficient in T cells ^40^ (**Figure 4A**). We confirmed the initial homing of cells to the lungs immediately after injection in both mouse strains (**Figure 4B**).

**Figure 4.**
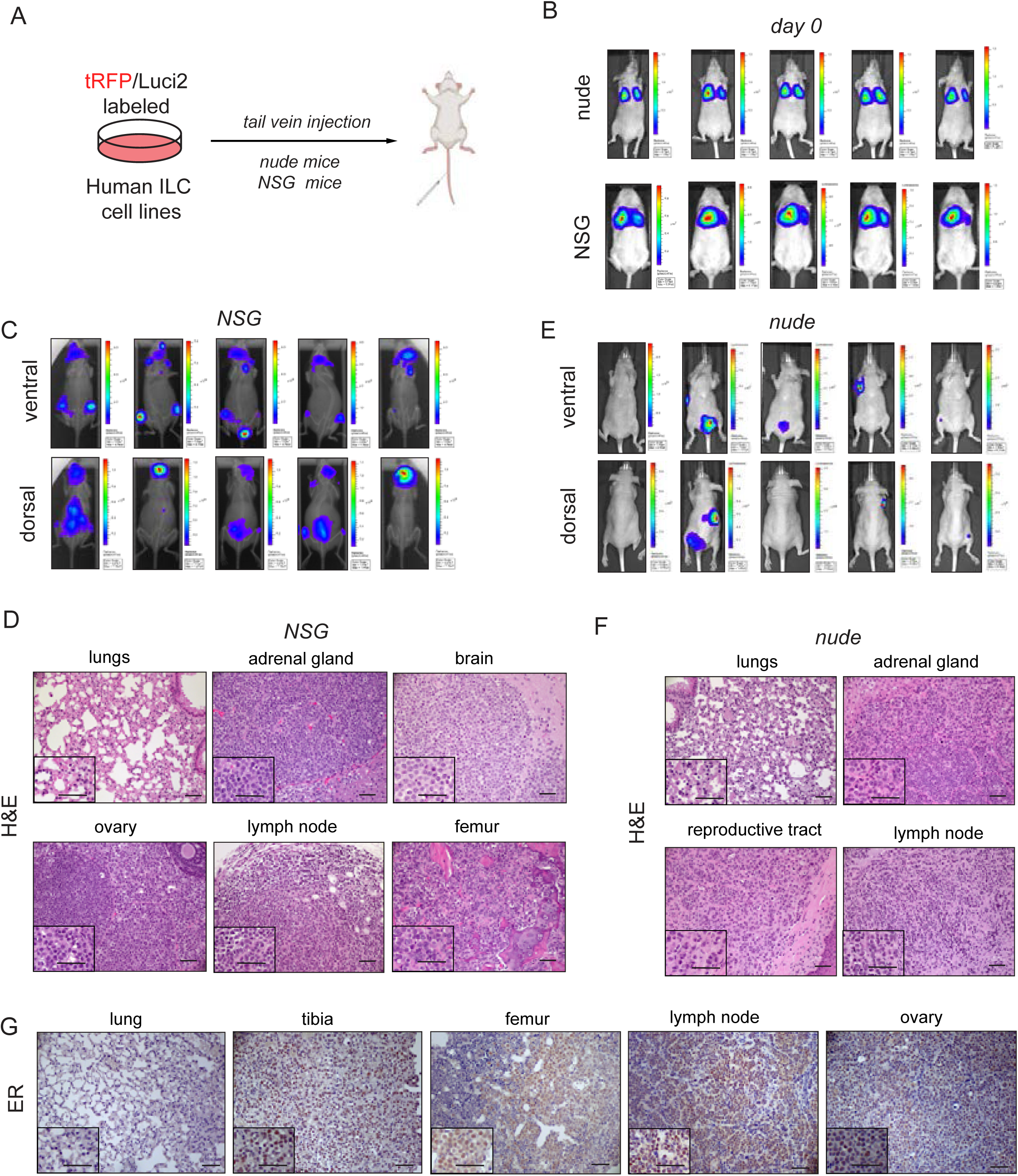
Human MDA-MB-134 tail vein xenografts give rise to experimental metastases to multiple sites. (A) Schematic of the experiment. (B) Ventral in vivo bioluminescent imaging of mice with MDA-MB-134 tail vein xenografts in nude (top) and NSG (bottom) mice immediately after injection. (C) Ventral (top) and dorsal (bottom) in vivo bioluminescent imaging of mice with MDA-MB-134 tail vein xenografts in NSG mice at end point. (D) H&E staining of experimental metastases from MDA-MB-134 tail vein xenografts to the indicated organs in NSG mice. (E) Ventral (top) and dorsal (bottom) in vivo bioluminescent imaging of mice with MDA-MB-134 tail vein xenografts in nude mice at end point. (F) H&E staining of experimental metastases from MDA-MB-134 tail vein xenografts to the indicated organs in nude mice. (G) ER staining of experimental metastases from MDA-MB-134 tail vein xenografts to the indicated organs in NSG mice. Lung images show absence of metastases. Insets show a small portion of the images at higher magnification. Scale bar: 40 μm.

In NSG mice, *in vivo* bioluminescent imaging revealed metastatic signal around one month post injection of MDA-MB-134 cells, which was localized to non-lung sites (**Figure 4C**). Histological evaluation by H&E staining confirmed the absence of lung colonization and the presence of extensive metastases to the adrenal gland, ovaries, lymph node, femur, and the brain (**Figure 4D**). As observed in the spontaneous metastases, these lesions were also positive for ER staining (**Figure 4G**). As expected, there was no signal in the lung, confirming absence of metastatic cells.

We next evaluated tail vein metastatic capacity in athymic nude mice. In nude mice, *in vivo* bioluminescence signal was observed around 1-3 months post injection of MDA-MB-134 cells and was also localized to non-lung sites (**Figure 4E**). *Ex vivo* fluorescence stereomicroscopy (**Figure S5A**) and histological evaluation by H&E staining (**Figure 4F**) confirmed the absence of lung colonization and the presence of MDA-MB-134 metastases to the adrenal gland, reproductive tract, bone and lymph node, although the metastatic burden was reduced compared to NSG mice. No metastatic signal was observed after tail vein injection of SUM44, MDA-MB-330 and BCK4 cells (**Figure S5B**; bottom and **Figure S5C**). In summary, tail vein injection of MDA-MB-134 cells results in wide-spread metastases primarily localized to non-lung sites, with more extensive metastatic spread observed in NSG compared to nude mice, likely reflecting the greater degree of immunodeficiency in this strain ^41^.

Given the extensive metastatic burden observed with ILC cell line xenografts, we asked if this dissemination could be captured by monitoring circulating cell-free DNA (cfDNA), which is under investigation in the clinic for diagnosis and prognosis ^42^. To this end, we generated MFP and tail vein xenografts with MDA-MB-134 cells and collected serum after the onset of metastasis (**Figure S6A**). MDA-MB-134 cells harbor a homozygous MAP2K4-S184L mutation (https://www.cbioportal.org/) and a heterozygous KRAS-G12R mutation ^32^. Wild-type and mutation-specific probes were initially validated by digital droplet PCR (ddPCR) on genomic DNA from MDA-MB-134 cells cultured *in vitro,* with MCF7 cells serving as a negative control (**Figure S6B** and **S6C**). ddPCR successfully detected the presence of *MAP2K4* and *KRAS* mutations in cfDNA extracted from the sera of mice bearing both MFP (**Figure S6D** and **S6E**) and tail vein (**Figure S6F** and **S6G**) xenografts, proving the feasibility of cfDNA tracking in these models.

In addition to the tail vein route, we also evaluated the potential of MDA-MB-134 and SUM44 cells to generate experimental metastases in athymic nude mice through the intracardiac route (**Figure 5A**), which introduces cells directly into the arterial blood supply and is commonly used to model bone metastasis ^43^. *In vivo* bioluminescent imaging confirmed the initial successful distribution of cells into systemic circulation immediately after injection (**Figure 5B**). Serial monitoring showed metastatic signal was detected starting at one-month post injection with MDA-MB-134 xenografts and localized extensively throughout the body (**Figure 5C**), while no signal was observed with SUM44 xenografts even at later time points (**Figure 5D**). Histological examination revealed metastases to the adrenal gland, femur, mandible, as well as to the eye (**Figure 5E**), with ER expression mostly notably in the femur (**Figure 5F**). All together, MDA-MB-134 xenografts exhibit a high potential of experimental metastasis via multiple routes in different immunocompromised backgrounds.

**Figure 5.**
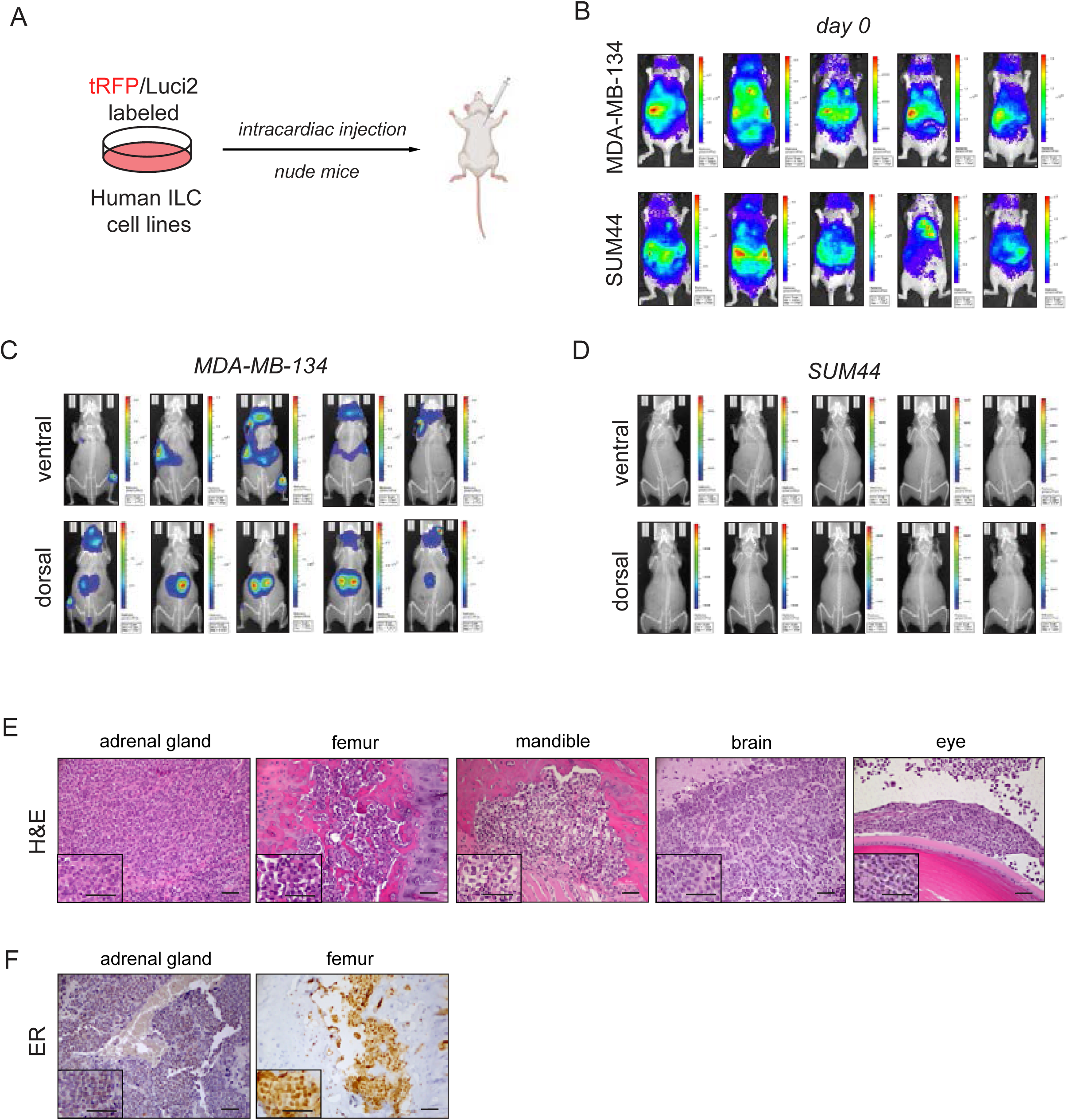
MDA-MB-134 but not SUM44PE intracardiac xenografts give rise to detectable metastases in nude mice. (A) Schematic of the experiment. (B) Ventral in vivo bioluminescent imaging of mice with MDA-MB-134 (top) and SUM44PE (bottom) intracardiac xenografts immediately after injection. (C-D) Ventral (top) and dorsal (bottom) in vivo bioluminescent imaging of mice with MDA-MB-134 (C) and SUM44PE (D) intracardiac xenografts at end point. SUM44PE images show absence of metastases. (E-F) H&E (E) and ER (F) staining of experimental metastases from MDA-MB-134 orthotopic xenografts to the indicated organs. Insets show a small portion of the images at higher magnification. Scale bar: 40 μm.

### Human ILC cell line orthotopic xenografts exhibit stromal cell recruitment and extracellular matrix deposition

Having characterized metastatic spread via multiple routes, we focused on the stromal composition and cellular microenvironment of the orthotopic mammary fat pad (MFP) xenografts. To understand the stromal composition of these MFP-derived primary tumors and metastatic lesions, we performed histological characterization of the tumor and metastatic microenvironments characterizing the extracellular matrix (ECM), as well as the presence of fibroblasts and neutrophils. Masson’s Trichrome staining revealed deposition of extracellular matrix in the form of collagen fibers, which was stronger in the primary tumors compared to the metastatic lesions (**Figures 6A-6C**). PDGFRβ staining confirmed presence of fibroblasts in both primary and metastatic tumors (**Figures 6A** and **6B**). These data show resemblance with stromal composition of transgenic ILC mouse models and human ILC tumors ^19^. NSG mice are deficient in mature lymphocytes and NK cells, but they retain neutrophils ^33^. Ly6G staining revealed varying degrees of neutrophil recruitment within the primary tumors (**Figure 6D**).

**Figure 6.**
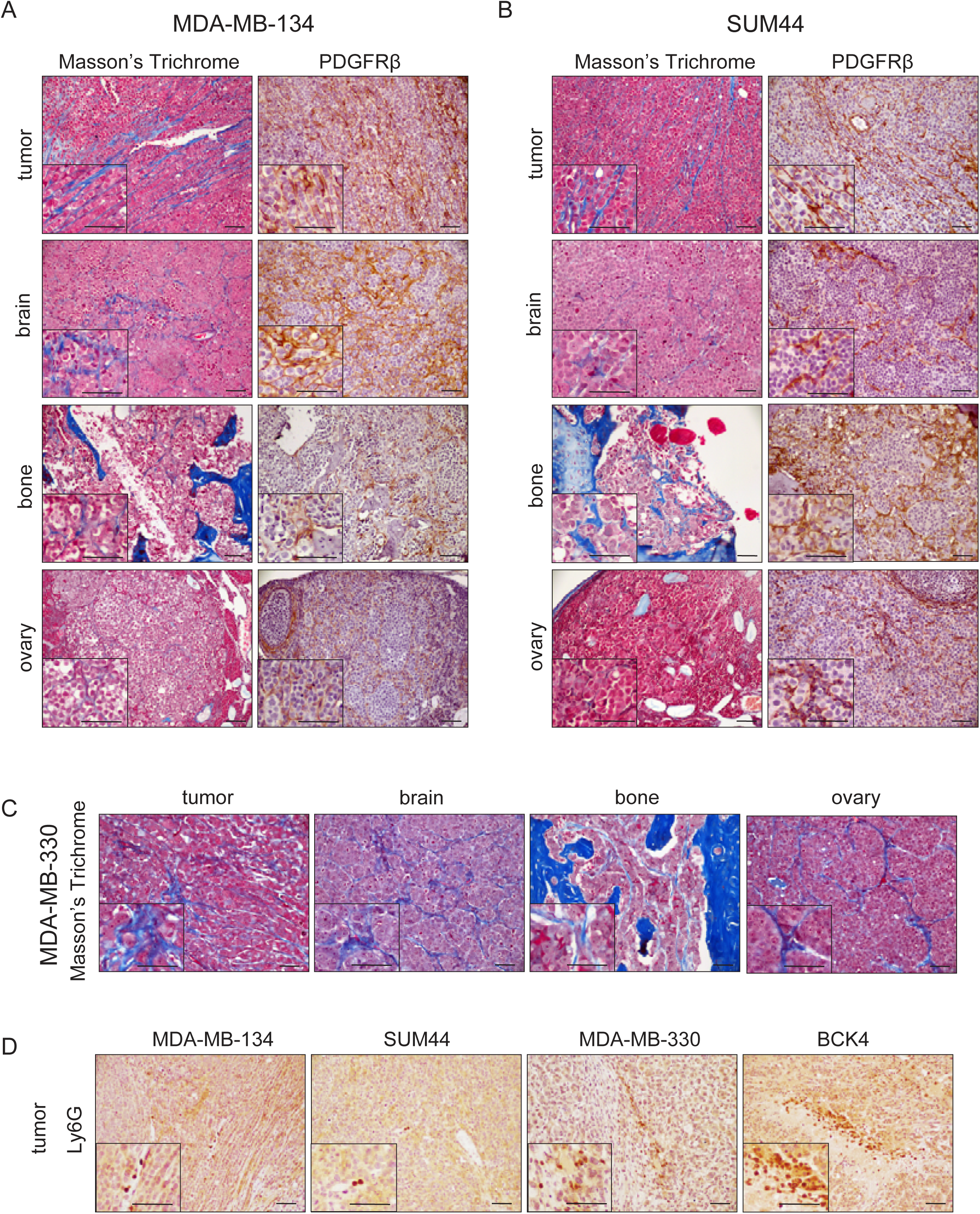
Human ILC cell line spontaneous metastases exhibit recruitment of stromal cells and collagen deposition. (A-B) Masson’s trichrome (left) and PDGFRβ (right) staining of primary tumors from MDA-MB-134 (A) and SUM44PE (B) orthotopic xenografts and spontaneous metastases to the indicated organs. (C) Masson’s trichrome staining of primary tumors from MDA-MB-330 orthotopic xenografts and spontaneous metastases to the indicated organs. (D) Ly6G staining of primary tumors from MDA-MB-134, SUM44PE, MDA-MB-330 and BCK4 orthotopic xenografts. Insets show a small portion of the images at higher magnification. Scale bar: 40 μm.

### ILC cell line xenografts are responsive to endocrine therapy

Having confirmed the expression of ER in the primary tumors and metastases of ILC cell line xenografts, we next sought out to evaluate response to endocrine therapy, which is the standard of care for patients with ER+ ILC ^9,12^. To this end, tRFP/luciferase-labeled MDA-MB-134 cells were used to generate orthotopic xenografts in NSG mice (**Figure 7A**). When the primary tumors reached 100-200 mm^3^ volume, the mice were randomized to either vehicle or the selective ER down-regulator (SERD) fulvestrant and monitored over time. Compared to the vehicle group, fulvestrant-treated mice displayed significant tumor growth inhibition (**Figures 7B-7D**). *In vivo* bioluminescent imaging captured significantly decreased burden from brain metastases (**Figures 7B**, **7E** and **7F**). *Ex vivo* bioluminescent imaging revealed less extensive colonization at additional metastatic sites in the fulvestrant group (**Figure 7F**), which also conferred a survival benefit to these mice (**Figure 7G**). IHC staining confirmed the downregulation of ER in the fulvestrant-treated cohort in both the primary tumor and at metastatic sites such as the brain and ovary (**Figure 7H**).

**Figure 7.**
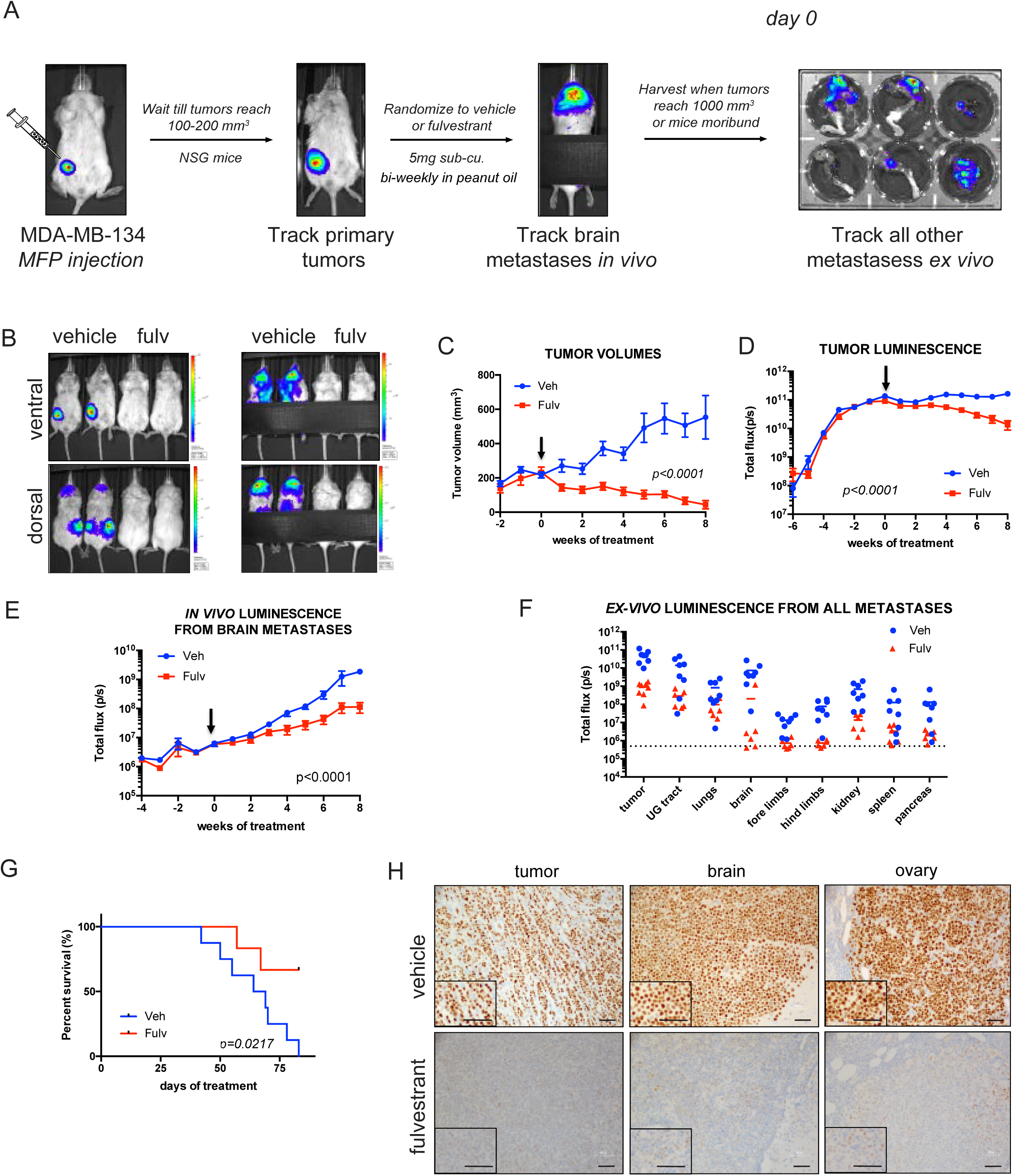
Tumors and metastases from MDA-MB-134 orthotopic xenografts are responsive to endocrine therapy. (A) Schematic of the experiment. (B) Ventral (top) and dorsal (bottom) in vivo bioluminescent imaging of mice with MDA-MB-134 orthotopic xenografts treated with vehicle or fulvestrant. In the images on the right, the lower bodies/primary tumors were covered to allow visualization of the spontaneous metastases to the upper body. (C-D) Volume growth (C) and in vivo bioluminescence (D) curves of primary tumors from MDA-MB-134 orthotopic xenografts treated with vehicle (blue) or fulvestrant (red). Note the saturation of the luciferase signal in the vehicle group in (D) around the start of treatment. (E-G) In vivo bioluminescence curves of brain metastases (E), ex vivo bioluminescence values of all experimental metastases (F) and survival curves (G) from mice with MDA-MB-134 orthotopic xenografts treated with vehicle (blue) or fulvestrant (red). Arrows indicate start of treatment. p-values for tumor volumes and total flux are from two-way ANOVA and for survival analysis from log-rank Mantel-Cox test. (H) ER staining of primary tumors and spontaneous metastases to the indicated sites from mice with MDA-MB-134 orthotopic xenografts treated with vehicle (blue) or fulvestrant (red). Insets show a small portion of the images at higher magnification. Scale bar: 40 μm.

In the above-described experiments, fulvestrant treatment was initiated before the onset of detectable metastases. To evaluate response to endocrine therapy in an established metastatic setting, we generated MDA-MB-134 xenografts in NSG mice via intracardiac injection, and randomized the mice to either vehicle or fulvestrant treatment after the detection of extensive metastatic burden (**Figure S7A**). Similar to spontaneous metastases from orthotopic xenografts, the experimental metastases from intracardiac xenografts at organs such as adrenal gland, bones and brain were significantly inhibited (**Figures S7B** and **S7C**) although there was no significant survival benefit in the fulvestrant cohort (**Figure S7D**). Collectively, these data demonstrate that ILC cell line xenografts can effectively model endocrine therapy, closely mirroring human ILC in the clinic.

### Transcriptional profiling captures characteristics of tumor growth, dissemination and microenvironment of human ILC xenografts

We performed transcriptional profiling of MDA-MB-134 xenografts to gain mechanistic insights into tumor growth and dissemination. We generated MDA-MB-134 orthotopic primary tumor xenografts and brain metastases in NSG mice and performed bulk RNA-Sequencing. We used XenofilteR to deconvolute the mouse and human RNA-Seq reads ^44^ yielding a “human reads dataset”. Comparison of the primary tumors versus the brain metastases via PCA showed general separate clustering between the two groups (**Figure 8A**). Consistent with the PCA, differential gene expression analysis revealed many transcripts (N=1019) (**Suppl Table S1**) significantly altered in the brain metastases compared to the primary tumors (**Figure 8B**). GSEA analysis revealed an enrichment in many pathways, including ER signaling (**Figure 8C**), as well as depletion of others including apoptosis/TP53, apical junctions and EMT. Given the unique ER-positivity of our ILC model, we followed up on the pathway analysis result and examined expression of genes within our recently reported “EstroGene” gene expression signatures ^45^ which showed robust increases in estrogen-induced genes such as RET and decreases in estrogen-repressed genes such as BLNK, indicating elevated estrogen receptor action in breast cancer brain metastases (**Figure 8D**).

**Figure 8:**
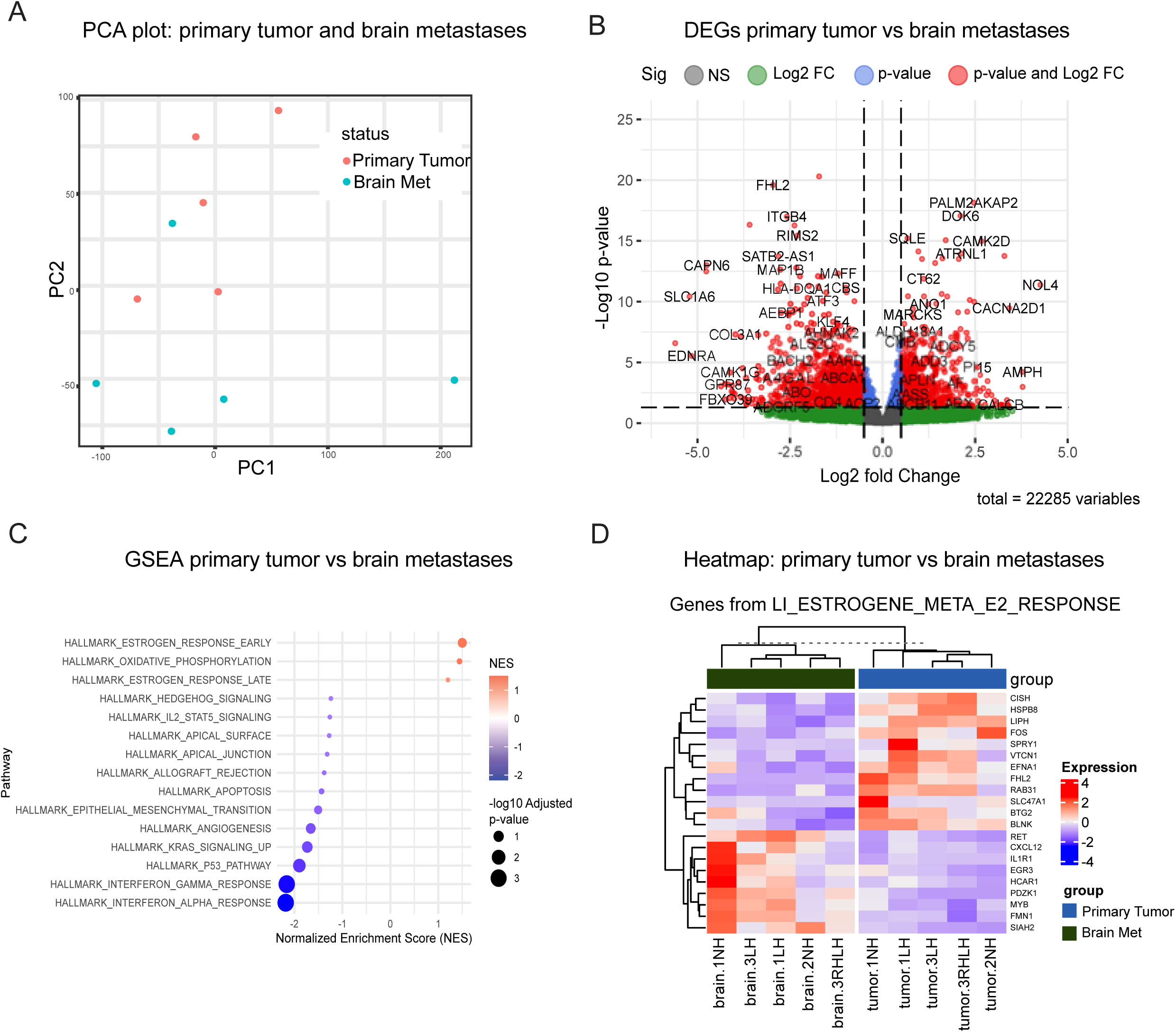
Transcriptional profiling of MDA-MB-134 orthotopic tumors and spontaneous brain metastases reveals shared and unique gene expression patterns. Analyses were performed using the human reads dataset obtained after XenofilteR deconvolution of mouse and human RNA-seq reads. (A) PCA plot of gene expression from MDA-MB-134 primary tumors and brain metastases. (B) Volcano plot showing differential gene expression between primary tumors and brain metastases. (C) GSEA analysis using the Hallmark gene set comparing primary tumors and brain metastases. (D) Heatmap showing significantly regulated genes from the Hallmark gene set LI_ESTROGENE_META_E2_RESPONSE_UP/LI_ESTROGENE_META_E2_RESPONSE_DN

## Discussion

ILC is a special histological subtype of breast cancer that is increasingly recognized for its unique features including its characteristic growth pattern, robust endocrine responsiveness and distinct metastatic profile ^1,6,12^. More and better in *vivo* models that can faithfully recapitulate clinical features of ILC are needed to increase our understanding of tumorigenesis and metastatic dissemination and enable development and testing of more effective therapeutic strategies. In this study, we generated orthotopic and experimental xenografts of human ILC cell lines in mice that mirror growth and metastatic behaviors seen in patients. The ability to detect disseminated disease by cfDNA tracking and the robust response observed to endocrine therapy support utility of these models as platforms for testing novel therapeutic hypotheses.

In orthotopic injection experiments, despite a generally slow *in vivo* tumor growth pattern, we detected spontaneous metastases to the brain, bones, ovaries, and other sites, closely mirroring clinical sites of dissemination of ILC ^1,12^ and capturing features such as invasion of leptomeningeal membranes ^36,37^ and bone erosion ^34,35^. Notably, similar ILC-specific metastatic dissemination patterns, including spread to bone, leptomeninges, and gynecologic tissues, have also been reported in MIND xenograft models supplemented with exogenous estradiol, further corroborating these as characteristic sites of ILC dissemination across multiple in vivo approaches ^46^. ERα, PR and Ecad/p120 expression was also consistent with *in vitro* observations ^26^ and studies on human tissues. ^1,12^. Stromal infiltration and extracellular matrix deposition were similar to that seen with other ILC mouse models ^19,20,23^ and patient samples ^1,6^.

MDA-MB-134 cells gave rise to more extensive metastases in NSG mice compared to nude mice. Interestingly, experimental lung metastases were not observed in either background, which was in stark contrast to the spontaneous lung metastases observed in NSG mice following mammary fat pad tumor growth. It is possible that primary tumors may educate the lung pre-metastatic niche to allow subsequent colonization, perhaps via exosomes secreted by the primary tumor, an education which may not occur in the tail vein injection model. MDA-MB-134 cells also gave rise to robust experimental metastases via the intracardiac route in nude mice, including colonization of the eye, reflective of orbital metastases seen in ILC patients ^14,47^. A more comprehensive analysis of SUM44, MDA-MB-330 and BCK4 cells via both routes and in both NSG and nude mice backgrounds could potentially reveal additional experimental metastases with these cell lines.

Transcriptomic profiling of orthotopic primary tumors and spontaneous metastases revealed several genes and pathways upregulated in brain metastasis, including ER signaling. Genes upregulated in brain metastases compared to primary tumors included the estrogen-regulated RET tyrosine kinase which we previously identified as one of the genes with the most recurrent expression gains in human breast cancer brain metastases ^48^. As ligands for RET such as Glial Cell Line-derived neurotrophic Factor (GDNF) show high expression in brain and have key functions for neuron survival, these findings warrant further interrogation in the context of ILC and may potentially reveal actionable therapeutic targets for metastatic control.

The retention of ERα expression in our human ILC cell line xenografts enabled us to test their response to endocrine therapy, which is the standard of care for patients with ILC ^9,12^. When mice were treated with the SERD fulvestrant, we observed a significant decrease in tumor and/or metastatic burden in both the orthotopic setting and in the established metastatic setting via the intracardiac route. These initial experiments were limited to MDA-MB-134 cells and future studies should investigate *in vivo* tumor/metastatic endocrine response of other human ILC cell lines. One limitation of our study was the use of different immunocompromised mouse strains to allow transplantation of human ILC cells.

While nude mice retain neutrophils but lack T cells, NSG mice have a more severe immunodeficiency affecting both lymphoid and myeloid lineages. This difference in immune competence likely influenced the metastatic efficiency observed, with NSG mice showing more extensive metastatic spread compared to nude mice for MDA-MB-134 tail vein xenografts, consistent with previous reports showing enhanced metastatic capacity in more immunocompromised backgrounds ^41^. These strain-specific differences highlight the important role of residual immune function in controlling metastatic dissemination and suggest that our findings reflect not only the intrinsic metastatic capacity of ILC cells but also their interaction with the host immune environment. While this enabled us to investigate human ILC biology to complement the learnings from transgenic mouse models ^19,20^, we were unable to fully capture the important role of the immune system in tumor growth and control. This could be better achieved by utilizing humanized mouse models in future studies ^49^. We also appreciate the development of other sophisticated models, such as patient derived organoids (PDO) and organs on a ChIP that will increasingly enable analysis of primary and advanced tumorigenesis. In sum, our study has added several new models and the associated knowledge to the growing number of *in vivo* resources available to the ILC research community.

## Acknowledgments

The authors thank Minji Chung and Weizhou Hou for critical technical support of the studies.

## Declaration of generative AI and AI-assisted technologies in the manuscript preparation process

During the preparation of this work the author(s) used Claude AI (Anthropic) and/or ChatGPT (OpenAI) in order to improve the clarity and readability of the English language. After using this tool/service, the author(s) reviewed and edited the content as needed and take(s) full responsibility for the content of the published article.

## Supplementary Figure Legends

**Figure S1. Orthotopic tumor growth of human ILC cell line xenografts in NSG mice.** (A) Individual tumor volumes from mice with BCK4, SUM44PE, MDA-MB-330, and MDA-MB-134 orthotopic xenografts. Red and blue colors indicate bilateral and unilateral injections into mammary fat pads. There was no significant difference in growth rates between bilateral and unilateral injections. Initially, 14 mice were injected for BCK4 and MDA-MB-134, and 15 mice were injected for SUM44 and MDA-MB-330. Final tumor growth data plotted: N=13 tumors for BCK4 and MDA-MB-134; N=14 tumors for MDA-MB-330; N=15 tumors for SUM44. Some mice did not develop measurable tumors and were excluded from analysis. (B) Xenograft tumor growth was modeled using a linear mixed-effects model with log-transformed tumor volume to compare the growth rate between cell lines. The the fitted straight of log-transformed tumor volume against day shows approximately linear trends for each cell line. Right panel shows formula and results which indicated significant differences between all cell line comparison except for MDA-MB-134 and SUM44 which exhibited similar growth rates. (C) Changepoints in tumor growth for BCK4 and MDA-MB-330 were calculated Using the R package strucchange. Prior to analysis, samples with essentially flat curves (very little growth) were excluded. In the left panel, the solid red vertical line indicates the estimated changepoint, representing the onset of detectable growth. The dashed lines show the confidence interval for this estimate. The adjusted p-value is also reported, providing statistical evidence for the significance of the detected growth onset. For BCK 4 as an example, most samples showed changepoints within a narrow window with an average changepoint day of 151, indicating a consistent onset of tumor growth across replicates and supporting the reliability of the detection. (D) Ventral in vivo bioluminescent imaging of mice with MDA-MB-134 orthotopic xenografts showing primary tumors. (E) Brightfield (left) and RFP (right) images of a representative MDA-MB-134 orthotopic tumor.

**Figure S2. MDA-MB-330 and BCK4 orthotopic xenografts give rise to ER+ metastases to clinically relevant anatomical sites.** (A-B) H&E, Ecad/p120, ER and/or PR staining of representative metastases from MDA-MB-330 (A) and BCK4 (B) orthotopic xenografts to the indicated organs. Insets show a small portion of the images at higher magnification. Scale bar: 40 μm.

**Figure S3. SUM44PE and MDA-MB-134 orthotopic xenografts give rise to ER+ metastases to additional clinically relevant anatomical sites.**

(A-B) H&E, Ecad/p120 and/or ER staining of representative metastases from MDA-MB-134 orthotopic xenografts to the spine (A) and uterine horn (B). Insets show a small portion of the images at higher magnification. Scale bar: 40 μm. (C) Human ER staining of representative metastases from MDA-MB-134 and SUM44PE to leptomeninges. Scale bar: 100 μm.

**Figure S4. MDA-MB-134 orthotopic xenografts rise to ER+ bone metastases featuring signs of bone erosion.** (A-B) Brightfield (top) and RFP (bottom) images (A) and H&E, Ecad/p120 and ER staining (B) of a representative spontaneous mandible metastasis from MDA-MB-134 orthotopic xenografts. (C-D) High resolution micro-CT image of the mandible metastasis in (A-B) in 3D MIP view (C) and 3D viewer (D). (E-F) High resolution micro-CT images of a representative femur metastasis (E) and femur metastasis (bone 1)/unaffected tibia (bone 2) (F) in 3D MIP view and as individual slices. White arrows indicate areas of bone erosion. Red arrow indicates unaffected bone.

**Figure S5. MDA-MB-134 but not SUM44PE, MDA-MB-330 or BCK4 tail vein xenografts give rise to detectable metastases in nude mice.** (A) Brightfield (top) and RFP (bottom) images of representative experimental metastasis from MDA-MB-134 tail vein xenografts in nude mice. (B) Ventral *in vivo* bioluminescent imaging of mice with SUM44PE, MDA-MB-330 and BCK4 tail vein xenografts in nude mice immediately after injection (top) and at end point (bottom). (C) H&E (top) and ER (bottom) staining of lungs from mice with SUM44PE, MDA-MB-330 and BCK4 tail vein xenografts in nude mice at end point showing absence of metastasis. Insets show a small portion of the images at higher magnification. Scale bar: 40 μm.

**Figure S6. Disseminated disease from MDA-MB-134 orthotopic and tail vein xenograft lesions can be tracked by monitoring cell free DNA in blood.** (A) Schematic of the experiment. (B-C) ddPCR analysis of MAP2K4 (B) and KRAS (C) mutation in MDA-MB-134 DNA (positive control) and MCF7 DNA/H2O (negative controls). (D-E) ddPCR analysis of MAP2K4 (D) and KRAS (E) mutation in serum cell free DNA from mice with MDA-MB-134 orthotopic xenografts/spontaneous metastases. (F-G) ddPCR analysis of MAP2K4 (F) and KRAS (G) mutation in serum cell free DNA from mice with MDA-MB-134 tail vein xenografts/experimental metastases. Serum cell free DNA from uninjected mice was used as negative control.

**Figure S7. Metastases from MDA-MB-134 intracardiac xenografts are responsive to endocrine therapy.** (A) Schematic of the experiment. (B) Ventral in vivo bioluminescent imaging of mice with MDA-MB-134 intracardiac xenografts treated with vehicle (left) or fulvestrant (right (C) Total in vivo bioluminescence curves of all detectable experimental metastases (C) and survival curves (D) from mice with MDA-MB-134 intracardiac xenografts treated with vehicle (blue) or fulvestrant (red).

