## Supplementary Figures for "Estrogen receptor-positive ILC cell line xenografts recapitulate metastatic dissemination and endocrine response of invasive lobular carcinoma"

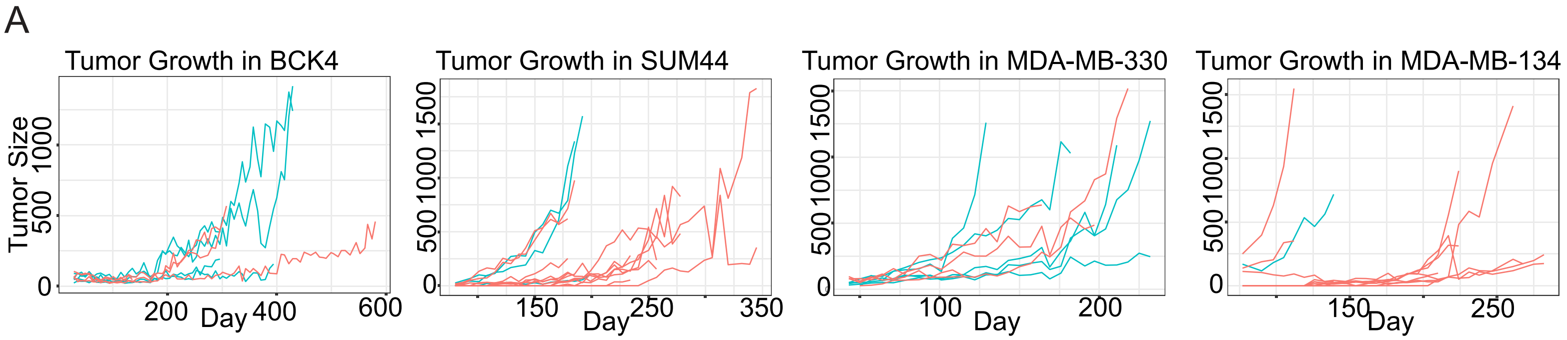


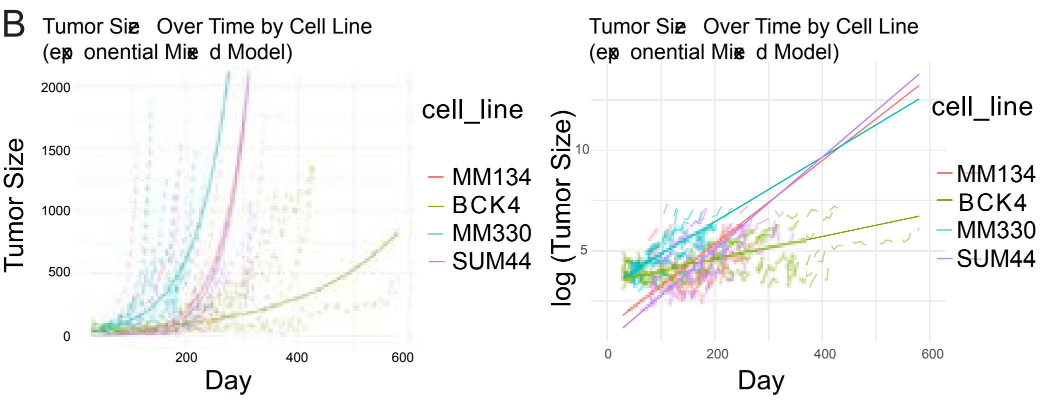


$$\log(TumorVolume_{ij} + 1) = \beta 0 + \beta 1 \cdot Day_{ij}$$
$$+ \Sigma(k=1..3)\ \gamma k \cdot I(cell\_line_{ij} = C_k)$$
$$+ \Sigma(k=1..3)\ \delta_k \cdot Day_{ij} \cdot I(cell\_line_{ij} = C_k)$$
$$+ b_{0j} + \varepsilon_{ij}$$

where:

$C_k \in$ {BCK4, MM330, SUM44}

I(.) is an indicator function (1 if true, 0 otherwise)

$\gamma_k$ = baseline offset for cell line $C_k$; $\delta_k$ = slope difference for cell line $C_k$

| contrast | estimate | SE | df | t.ratio | p.value |
|---|---|---|---|---|---|
| MM134 BCK4 | 0.02822 | 0.00112 | 1333 | 25.268 | <.0001 |
| MM134 MM330 | 0.01777 | 0.00145 | 1331 | 12.218 | <.0001 |
| MM134 SUM44 | 0.00166 | 0.00138 | 1332 | 1.279 | 0.5765 |
| BCK4 MM330 | -0.01845 | 0.00192 | 1328 | -19.72 | <.0001 |
| BCK4 SUM44 | -0.0256 | 0.00079 | 1327 | -33.618 | <.0001 |
| MM330 SUM44 | -0.01611 | 0.00122 | 1328 | -13.186 | <.0001 |

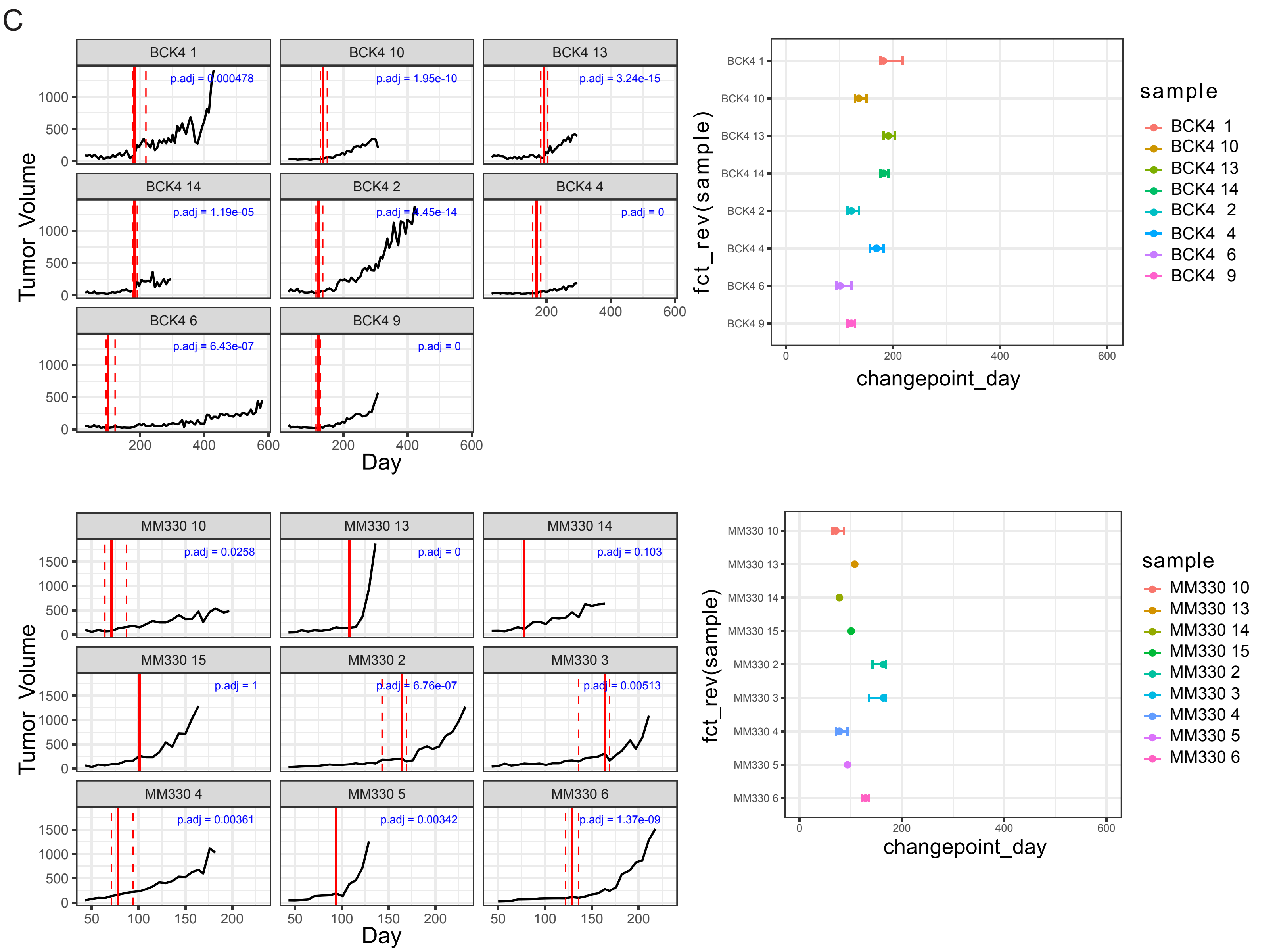

D

***in vivo***

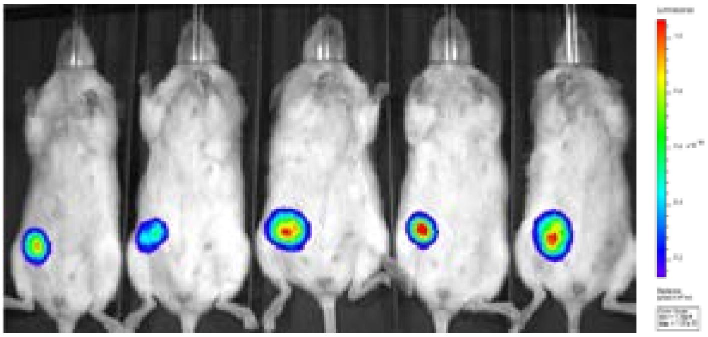

E

***ex vivo***

Brightfield RFP

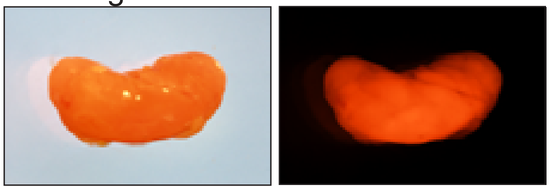

**Fig S1. Orthotopic tumor growth of human ILC cell line xenografts in NSG mice.**

Fig.S2

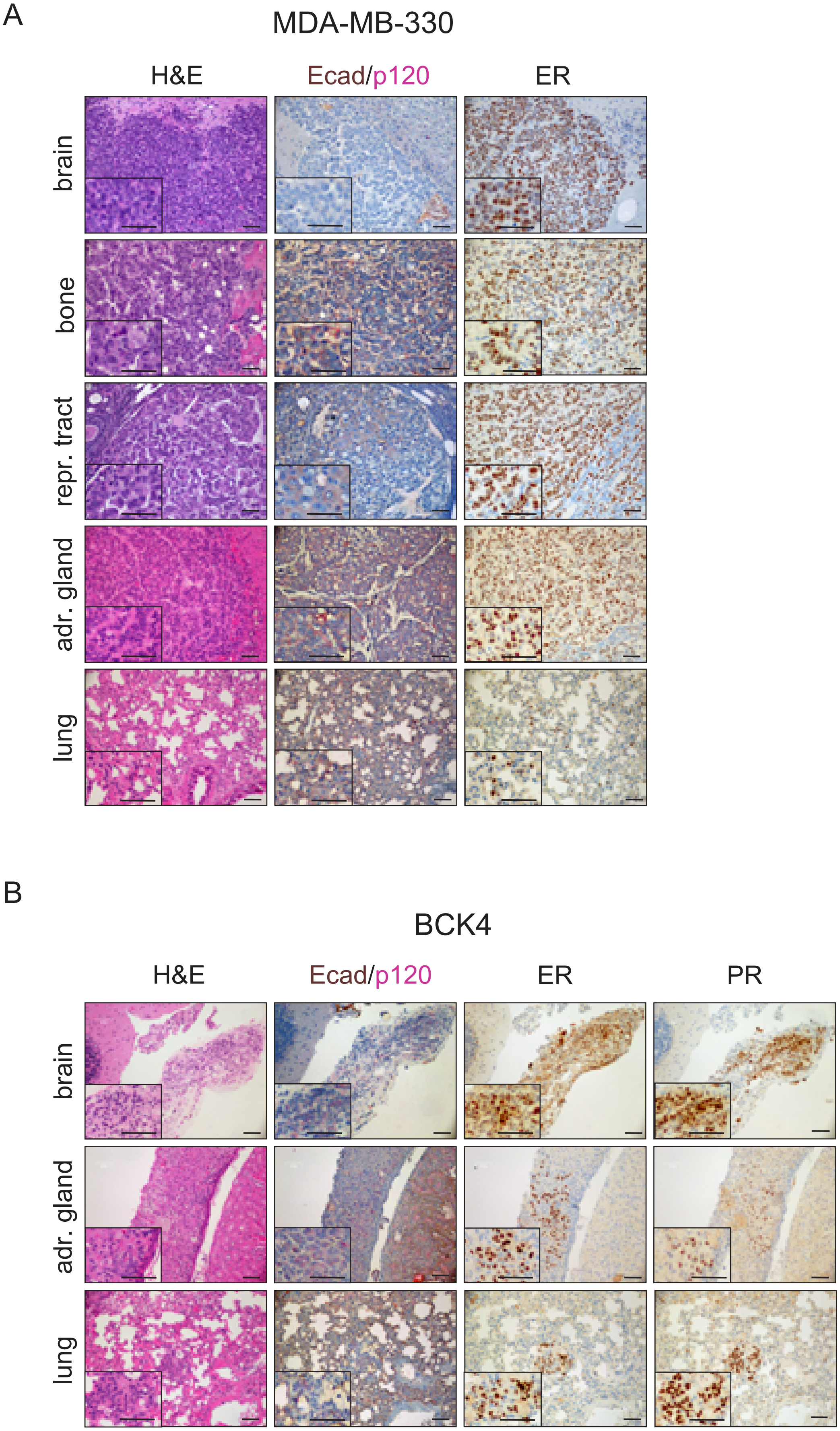


**Fig S2. MDA-MB-330 and BCK4 orthotopic xenografts give rise to ER+ metastases to clinically relevant anatomical sites.**

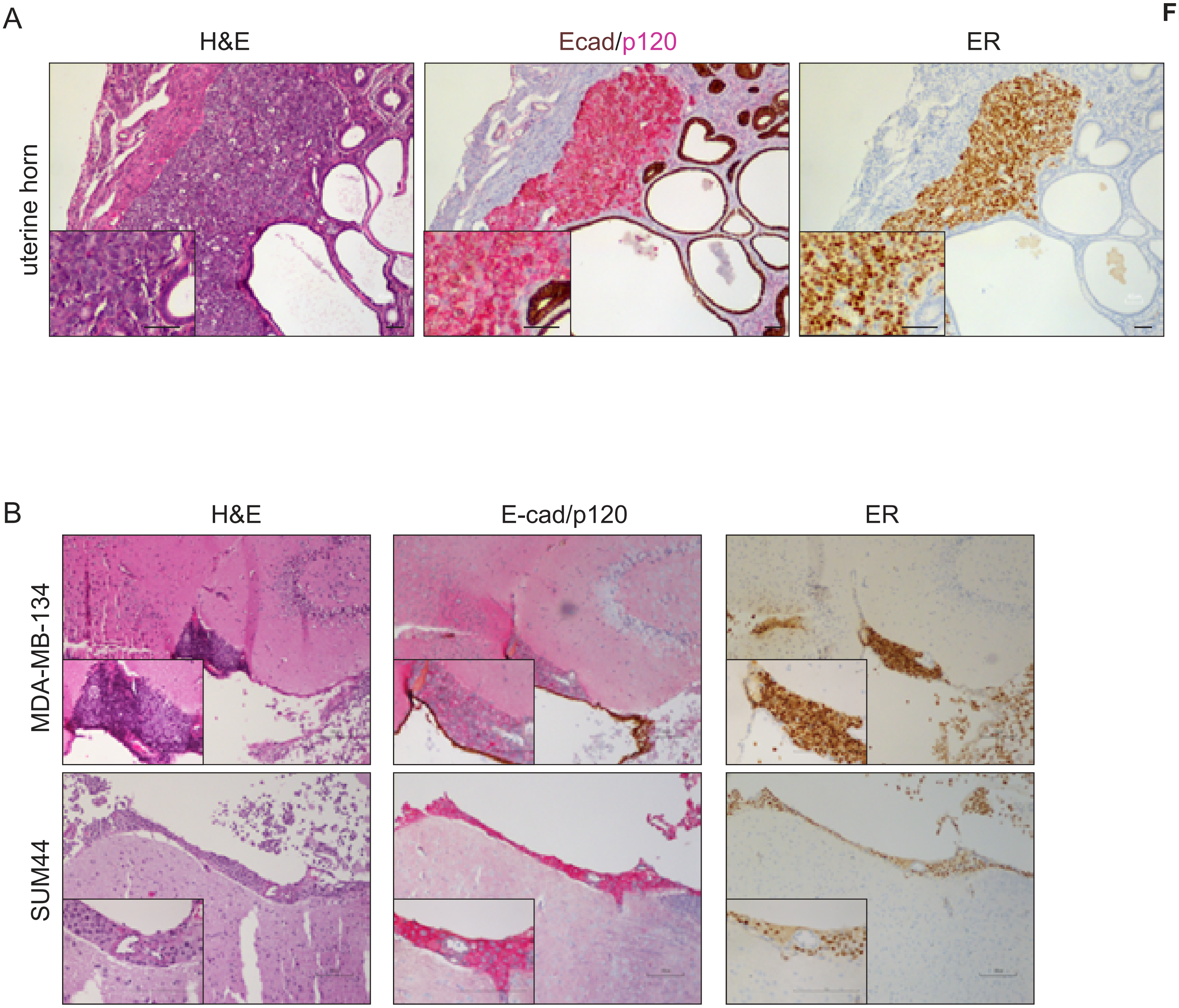


**Fig S3. SUM44PE and MDA-MB-134 orthotopic xenografts give rise to ER+ metastases to additional clinically relevant anatomical sites.**

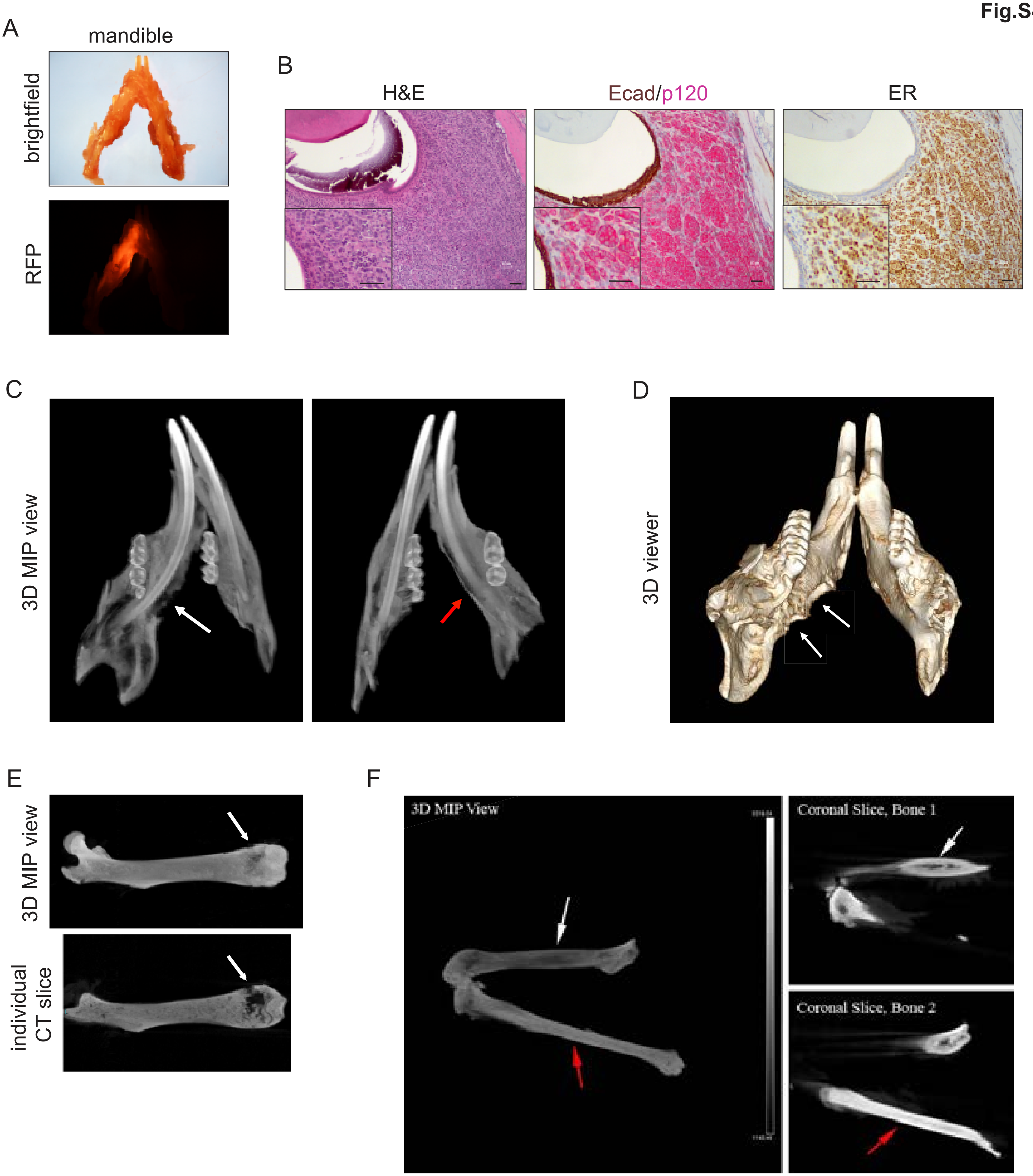


**Fig S4. MDA-MB-134 orthotopic xenografts give rise to ER+ bone metastases featuring signs of bone erosion.**

Fig.S5

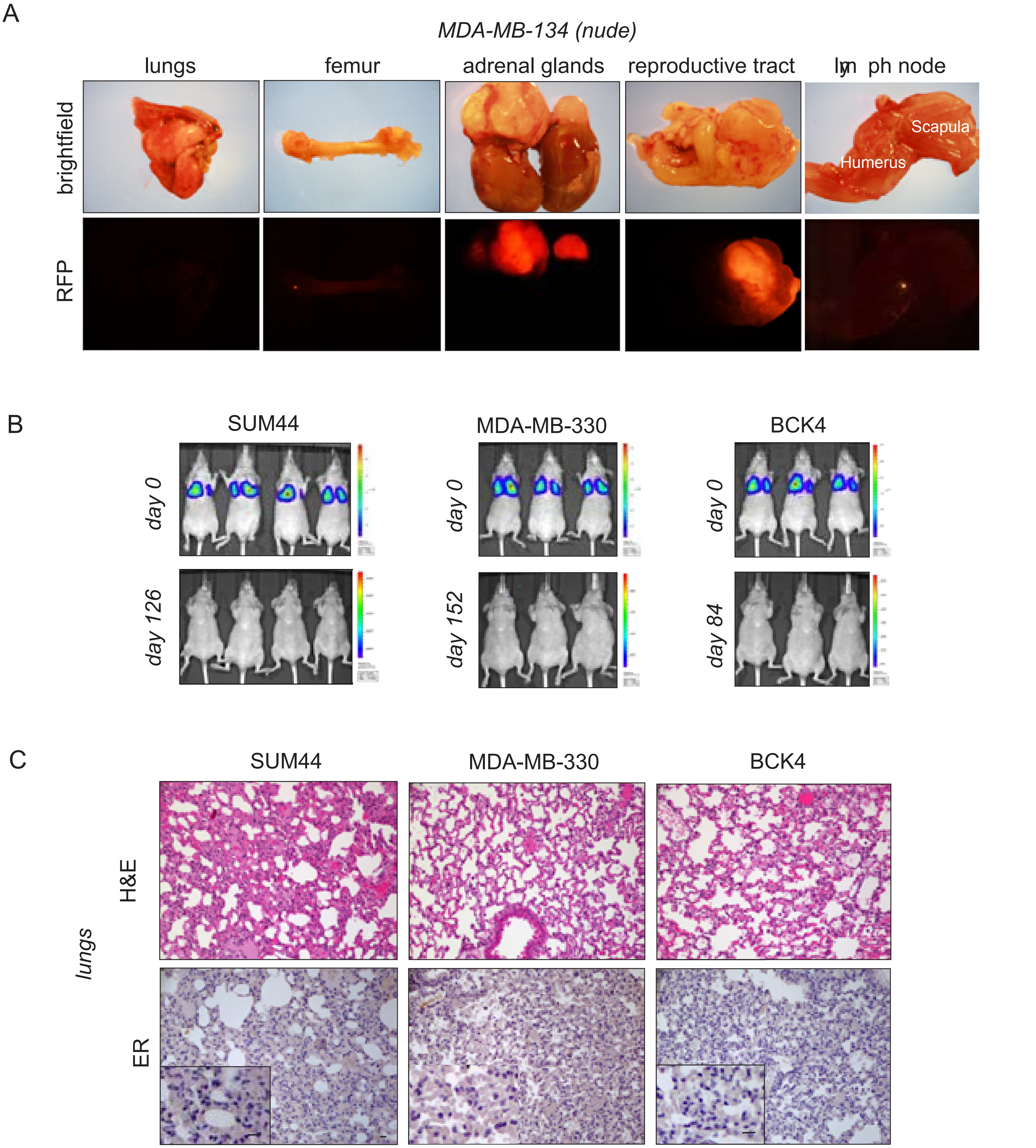


**Fig S5. MDA-MB-134 but not SUM44PE, MDA-MB-330 or BCK4 tail vein xenografts give rise to detectable metastases in nude mice.**

Fig. S6

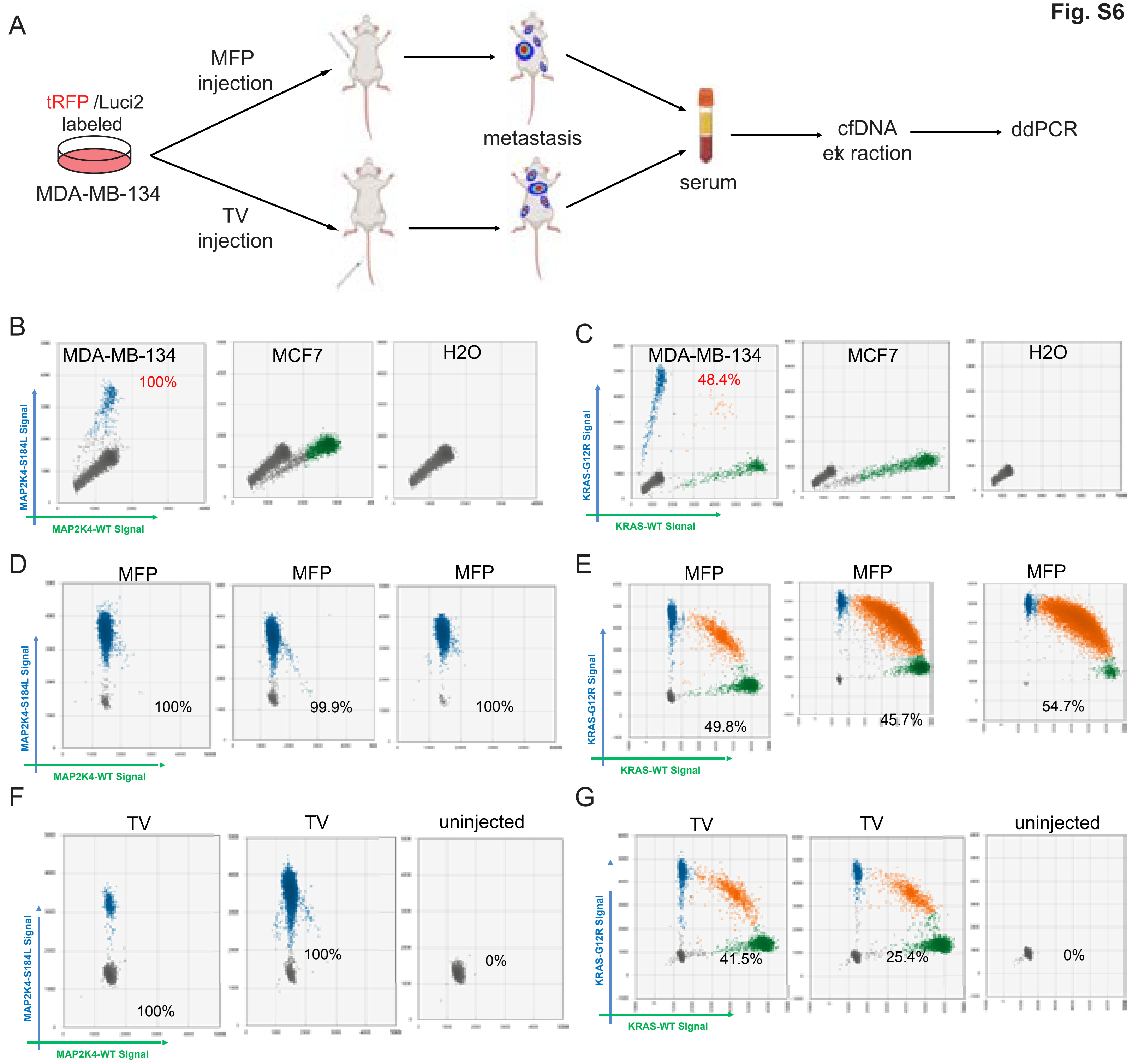


**Fig S6. Disseminated disease from MDA-MB-134 orthotopic and tail vein xenograft lesions can be tracked by monitoring cell free DNA in blood.**

Fig.S7

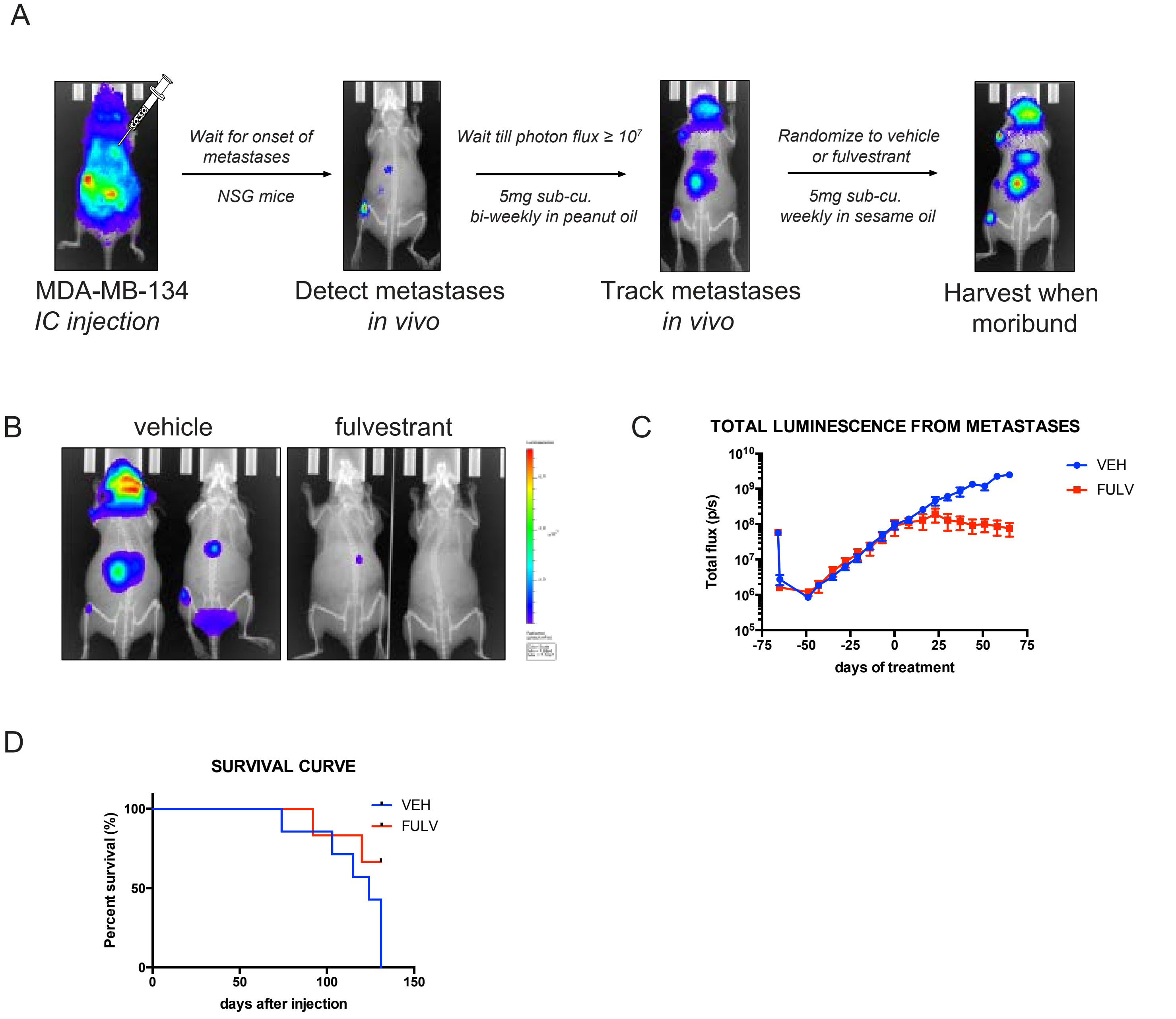


**Fig S7. Metastases from MDA-MB-134 intracardiac xenografts are responsive to endocrine therapy.**
